# Reduced Thalamic Excitation to Motor Cortical Pyramidal Tract Neurons in a Mouse Model of Parkinsonism

**DOI:** 10.1101/2022.09.24.509340

**Authors:** Liqiang Chen, Samuel Daniels, Rachel Dvorak, Hong-Yuan Chu

## Abstract

Degeneration of midbrain dopaminergic (DA) neurons causes a reduced motor output from the primary motor cortex (M1), underlying the motor symptoms of Parkinson’s disease (PD). However, cellular and circuitry mechanisms of M1 dysfunction in PD remain undefined. Using multidisciplinary approaches, we found that DA degeneration induces cell-subtype- and inputs-specific reduction of thalamic excitation to M1 pyramidal tract (PT) neurons. Physiological and anatomical analyses suggest that DA degeneration induces a loss of thalamocortical synapses to M1 PT neurons, resulting in an impaired thalamic driving of their activities. Moreover, we showed that the decreased thalamocortical connectivity are mediated by an excessive activation of NMDA receptors of M1 PT neurons. Further, the decreased thalamocortical transmission in parkinsonism can be rescued by chemogenetically suppressing basal ganglia outputs. Together, our data suggest that the reduced motor cortical outputs in parkinsonism are not only an immediate consequence of basal ganglia inhibition but also involves specific local circuitry adaptations within M1. This study reveals novel insight in the pathophysiology of parkinsonian motor deficits.

## Introduction

Degeneration of dopaminergic (DA) neurons in the substantia nigra pars compacta (SNc) is one of featured pathology of Parkinson’s disease (PD), which is causally linked with the parkinsonian motor deficits. Following the loss of the SNc DA neurons, both magnitude and timing of GABAergic outputs from the basal ganglia to downstream motor regions are robustly altered, which cause an abnormal suppression of the initiation and execution of movements in PD (Albin et al., 1989; Galvan and Wichmann, 2008; McGregor and Nelson, 2019). The thalamocortical network integrates basal ganglia information and sends motor commands to the motor areas in the brain stem and the spinal cord for movement execution, placing it an ideal position to mediate many aspects of motor symptoms of PD, particularly those related to skilled motor activities.

The primary motor cortex (M1) plays a critical role in the acquisition and execution of skilled motor activity (Georgopoulos and Carpenter, 2015; Guo et al., 2015a; Shenoy et al., 2012). M1 sends temporal patterns of activity for the generation and execution of movements and such network dynamics depends on both intrinsic mechanisms and synaptic inputs (e.g., thalamic excitation) (Sauerbrei et al., 2020; Shenoy et al., 2012). In parkinsonian state, the pathological signals of the basal ganglia are predicted to decrease M1 output and/or disrupt its network dynamics, contributing to the manifestation of the hypokinetic symptoms in PD (Aeed et al., 2021; Brazhnik et al., 2016; Hyland et al., 2019; McGregor and Nelson, 2019). However, our understanding of M1 pathophysiology in parkinsonism remains incomplete.

A large body of evidence in literature suggests M1 circuitry dysfunction in parkinsonism. Degeneration of SNc DA neurons impairs motor skills and decreases forelimb representation in the M1 of rodents (Brown et al., 2009; Plowman et al., 2011; Viaro et al., 2011), which perhaps involve intrinsic and synaptic adaptations in M1 associated with DA depletion (Aeed et al., 2021; Cousineau et al., 2020; Guo et al., 2015b; Villalba et al., 2021). Consistently, M1 pyramidal neurons exhibit enhanced bursting pattern of activity and synchronization following chronic DA degeneration (Aeed et al., 2021; Goldberg et al., 2002; Pasquereau and Turner, 2011; Pasquereau et al., 2016). For example, in MPTP-treated nonhuman primates (NHP), while the layer 5 pyramidal tract (PT) neurons exhibit a decreased rate and enhanced bursting pattern of activity, the intratalencephalic (IT) neurons are largely intact (Pasquereau and Turner, 2011). In addition, both electrophysiology and optical imaging studies suggest an impaired timing of PT neuronal activation and the execution of movement in parkinsonian NHP and rodents (Aeed et al., 2021; Pasquereau et al., 2016). In agreement with the earlier studies, our recent work reported that the PT, but not the IT, neurons in M1 show selective reduction in their intrinsic excitability, and that such intrinsic adaptations involve a combination of several ionic conductance (Chen et al., 2021). Therefore, in addition to the acute effects of the abnormal basal ganglia GABAergic inhibition, M1 circuits exhibit local adaptations at the cellular and synaptic levels, which can be important aspects of the pathophysiology of motor deficit in PD.

Neural circuits are equipped with a variety of homeostatic synaptic plasticity mechanisms to stabilize network function in face of perturbations (Turrigiano, 2011). For example, our recent work demonstrated several synaptic and intrinsic adaptations in the basal ganglia associated with pathological neuronal activity and DA degeneration (Chu et al., 2017; McIver et al., 2019). Based on the abnormal rate and pattern of M1 neuronal activity in parkinsonism, we hypothesized that loss of DA neurons induces cell-subtype- and input-specific synaptic adaptations of the glutamatergic inputs of M1 and that such synaptic adaptations contribute to the reduced cortical output in parkinsonism. A multidisciplinary approach was used to test these hypotheses in a well-established neurotoxin-based mouse model of parkinsonism.

## Results

Mice received either 6-OHDA injections into the medial forebrain bundle (MFB) to induce the degeneration of SNc DA neurons (“6-OHDA mice” hereafter), or vehicle injections into the MFB as controls (“controls” hereafter). All physiology and anatomical studies were conducted between 3–5 weeks post-injections. To confirm a complete degeneration of nigrostriatal DA pathway in 6-OHDA mice, all animals were subject to behavioral tests at 3 weeks post-surgery, prior to electrophysiology and anatomy studies (Chen et al., 2021). Relative to controls, 6-OHDA mice showed a significant reduction of locomotor activity in open field test (distance travelled in 10 min, control = 39 [33–48] m, n = 51 mice; 6-OHDA = 21 [15–27] m, n = 52 mice; *p* < 0.0001, MWU test). Moreover, 6-OHDA mice also exhibited asymmetric forelimb use in cylinder test (% of ipsilateral forelimb use, controls = 50 [49–52]%, n = 51 mice; 6-OHDA = 85 [79– 93]%, n = 52 mice; *p* < 0.0001, MWU test). *Post hoc* immunostaining of striatal TH was conducted, which showed a > 80% loss of nigrostriatal axon terminals in the lesioned hemisphere of 6-OHDA mice (%striatal TH_ipsilateral /contralateral TH_, control = 100 [97–102]%, n = 51 mice; 6-OHDA = 11 [7–16]%, n = 52 mice; *p* < 0.0001, MWU test).

### SNc DA degeneration induces cell-subtype- and input-specific alterations in M1 circuits

To interrogate synaptic properties of thalamocortical inputs to projection-defined subtypes of M1 layer 5 (L5) pyramidal neurons in controls and 6-OHDA mice, (1) AAV9-hSyn-ChR2(H134R)-eYFP were injected into the motor thalamus (centering at the ventromedial (VM) subregion of thalamus, **Fig. 1B**); (2) Retrobeads were stereotaxically injected into the ipsilateral pontine nuclei and contralateral striatum to label PT and IT neurons, respectively (**Figure 1A**). Three weeks post-injection, intense eYFP labeled fibers could be observed, particularly in the layers I and V of M1 (**Figure 1B**). Synaptic strength of thalamic inputs to PT and IT neurons were compared using *ex vivo* electrophysiology and optogenetics in slices from controls and 6-OHDA mice.

**Figure 1.**
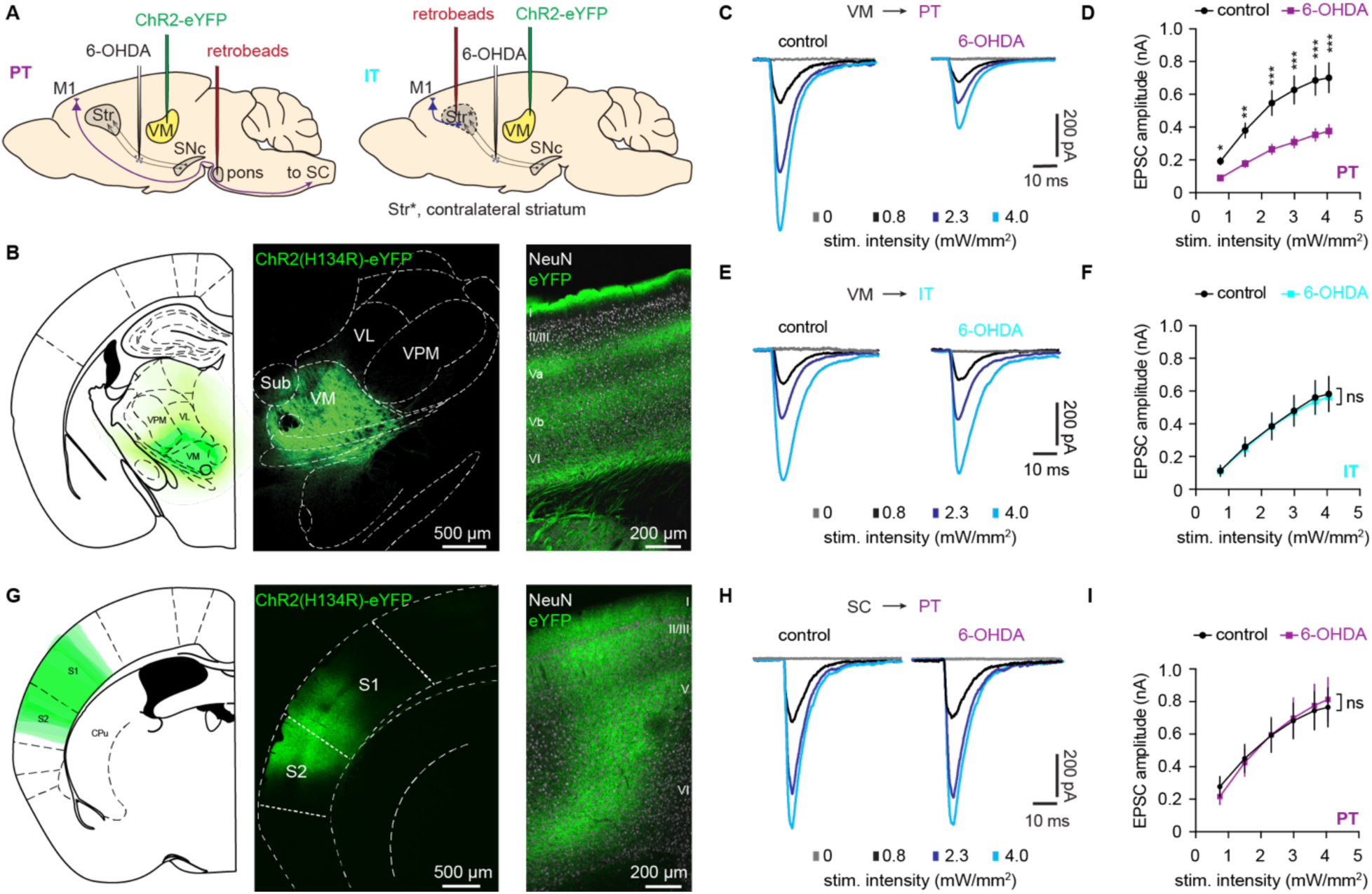
SNc DA degeneration induces cell-subtype- and input-specific alterations in M1. **A**) Schematic of overall strategies to retrogradely label M1 pyramidal neurons based on their projections and AAV-mediated optogenetics to selectively activate thalamocortical transmission arising from the ventromedial (VM) subregions of the motor thalamus of both controls and 6-OHDA mice. **B**) AAV-infected thalamic regions were overlaid to show the center of AAV injections across animals (left) and representative confocal images showing AAV9-ChR2(H134R)-eYFP infusion site in the motor thalamus that was centered at the VM subregion middle), and ChR2(H134R)-eYFP-expressing thalamic axon terminals in the M1, which concentrate in the layers I and V (right). **C-D**) Representative traces of optogenetically-evoked EPSCs across different stimulation intensities in PT neurons from both controls and 6-OHDA mice (**C**) and the summarized results (**D**). **E-F**) Representative traces of optogenetically-evoked EPSCs across different stimulation intensities in IT neurons from both controls and 6-OHDA mice (**E**) and the summarized results (**F**). **G**) AAV-infected cortical regions were overlaid to show the center of AAV injections across animals (left) and representative confocal images showing AAV9-ChR2(H134R)-eYFP infusion site in the motor thalamus that was centered at the sensory cortical areas (middle), and ChR2(H134R)-eYFP-expressing cortical axon terminals in the M1 right). **H-I**) Representative traces of optogenetically-evoked EPSCs arising from the SC across different stimulation intensities in M1 PT neurons from both controls and 6-OHDA mice (**H**) and the summarized results (**I**). *, *p* < 0.05, **, *p* < 0.01, ***, *p* < 0.001, ns, not significant.

Whole-cell voltage-clamp recordings of excitatory postsynaptic currents (EPSCs) were conducted in the presence of TTX (1 μM) and 4-AP (100 μM) to prevent the recruitment of polysynaptic responses and facilitate presynaptic glutamate release from thalamic axon terminals (Hooks et al., 2013; Petreanu et al., 2007). The amplitude of optogenetically-evoked thalamic EPSCs (oEPSCs) in M1 PT neurons was greatly reduced in slices from 6-OHDA mice relative to those from controls across a range of stimulation intensities (*p* < 0.0001, 2-way ANOVA followed by Sidak’s tests, **Figure 1C, D**). In contrast, the amplitude of oEPSCs in the IT neurons was unaltered between slices from 6-OHDA mice and controls (*p* > 0.05, 2-way ANOVA, **Figure 1E, F**). These data suggest that chronic SNc DA degeneration selectively decreases synaptic strength of the motor thalamic excitation to M1 PT neurons.

Cerebral cortical neurons are equipped with a battery of homeostatic mechanisms to scale-up or -down synaptic strength to compensate for external perturbations (Turrigiano, 2011). Thus, we examined whether loss of SNc DA neurons decreases global excitation to PT neurons through homeostatic plasticity processes, or selectively decreases thalamic inputs? To answer this question, AAV-hsyn-ChR2(H134R)-eYFP were injected into the sensory cortical (SC) regions (**Figure 1G**) and retrobeads were injected into pons of 6-OHDA mice and controls. Next, we compared the connection strength of sensory cortical inputs to M1 PT neurons of 6-OHDA mice and controls. The amplitude of oEPSCs from SC to PT neurons (SC-PT) was not altered in slices from 6-OHDA mice relative to controls (**Figure 1H, I**). These data suggest that DA degeneration likely selectively decrease thalamic excitation to M1 PT neurons. Together, chronic DA degeneration leads to cell-subtype- and input-specific adaptions in M1 local circuits.

### SNc DA degeneration decreases the number of functional thalamic inputs to M1 PT neurons

To gain mechanistic understanding of synaptic adaptations of M1 circuits following the loss of SNc DA neurons, we assessed the quantal properties of glutamatergic neurotransmission to PT and IT neurons in M1. Replacing extracellular Ca^2+^ with Sr^2+^ (2 mM), we found a significantly reduced frequency, but an unaltered amplitude, of optogenetically-evoked, Sr^2+^-induced thalamic asynchronous EPSCs (Sr^2+^-EPSCs) in the PT neurons from 6-OHDA mice relative to those from controls (frequency of Sr^2+^-EPSCs, controls = 34 [15.5–52] Hz, n = 20 neurons/3 mice; 6-OHDA = 20 [12–30] Hz, n = 20 neurons/4 mice; *p* = 0.02, MWU; amplitude of Sr^2+^-EPSCs, controls = 7.95 [6.3– 13.2] pA, n = 20 neurons/3 mice; 6-OHDA = 10.2 [8.4–12.8] pA, n = 20 neurons/4 mice; *p* = 0.33, MWU, **Figure 2A-C**). In contrast, neither the frequency nor the amplitude of thalamic Sr^2+^-EPSCs in the IT neurons was altered in 6-OHDA mice relative to controls (frequency of Sr^2+^-EPSCs, controls = 34.8 [15.7–50.5] Hz, n = 22 neurons/3 mice; 6-OHDA = 38.4 [24.9–47.6] Hz, n = 22 neurons/3 mice; *p* = 0.7, MWU; amplitude of Sr^2+^-EPSCs, control = 12.6 [10.5–14.4] pA; 6-OHDA = 11.8 [10.8–12.4] pA; n = 22 neurons/3 mice; *p* = 0.34, MWU; **Figure 2D-F**). Consistent with the intact synaptic strength of sensory cortical inputs to PT neurons (**Figure 1G-I**), the frequency and amplitude of Sr^2+^-EPSCs at SC–PT synapses were unaltered between groups (frequency of Sr^2+^-EPSCs, controls = 48 [29–68] Hz, n = 20 neurons/3 mice; 6-OHDA = 31.2 [26–52] Hz, n = 19 neurons/3 mice; *p* = 0.6, MWU; amplitude of Sr^2+^-EPSCs, control = 17 [16–20.8] pA, n = 20 neurons/3 mice; 6-OHDA = 16.5 [14.4–19.3] pA, n = 19 neurons/3 mice; *p* = 0.5, MWU; **Figure 2G-I**). These data further support that SNc DA degeneration selectively affect thalamic projection to PT neurons, largely due to fewer functional VM-PT inputs.

**Figure 2.**
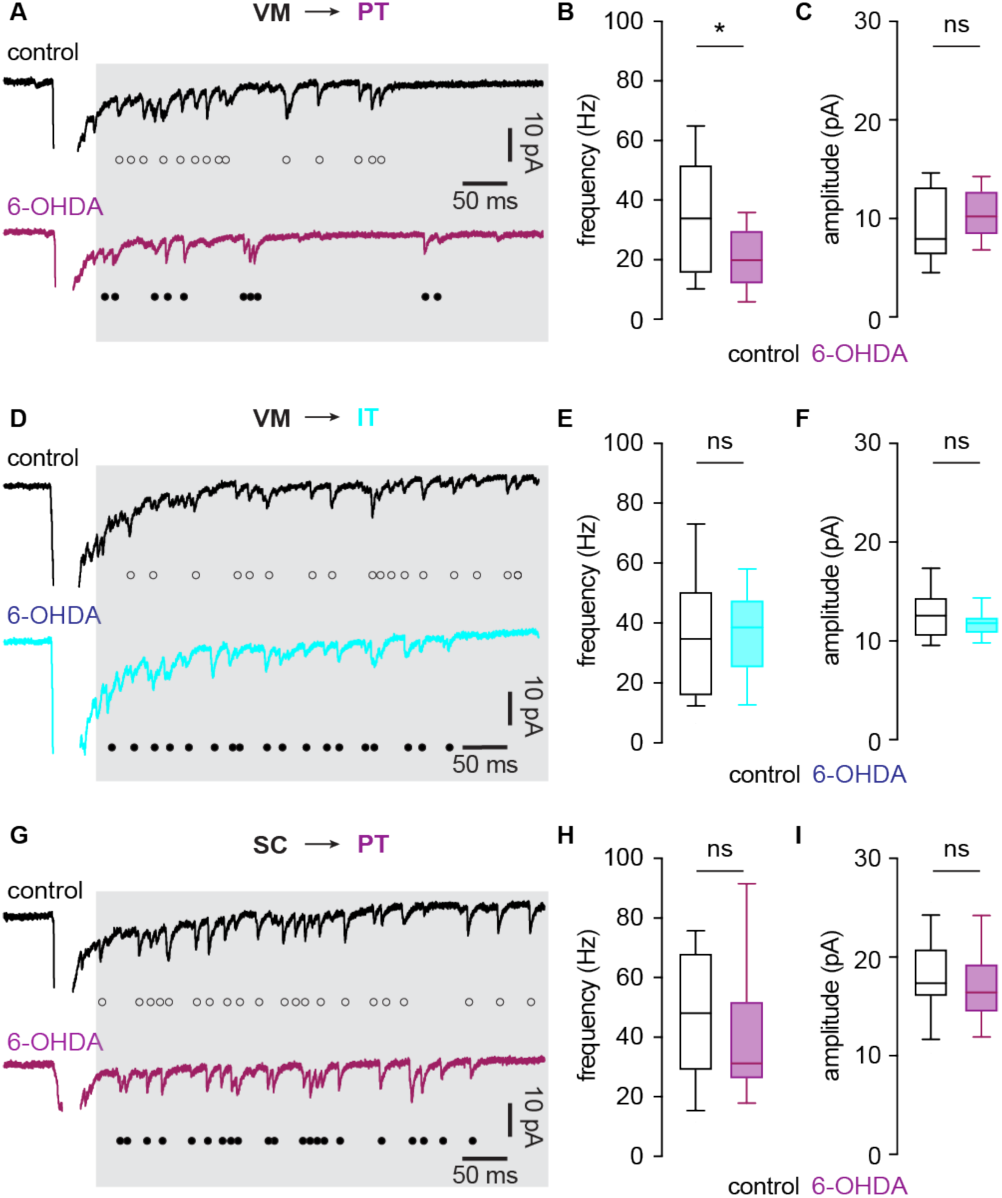
SNc DA degeneration decreases the number of functional thalamic inputs to M1 PT neurons. **A**) Representative traces showing optogenetically-evoked, Sr^2+^-induced asynchronous thalamocortical EPSCs of PT neurons from controls and 6-OHDA mice. Open and filled circle indicate individual events of asynchronous EPSCs. **B-C**) Box plots showing a reduced frequency (B) and unaltered amplitude (C) of Sr^2+^-EPSCs at thalamic inputs to M1 PT neurons from 6-OHDA mice relative to controls. **D**) Representative traces showing optogenetically-evoked, Sr^2+^-induced quantal EPSCs of thalamic inputs to IT neurons from controls and 6-OHDA mice. Open and filled circle indicate individual events of asynchronous EPSCs. **E-F**) Box plots showing unaltered frequency (E) and amplitude (F) of Sr^2+^-EPSCs at thalamic inputs to M1 IT neurons between groups. **G**) Representative traces showing optogenetically-evoked, Sr^2+^-induced quantal EPSCs of sensory cortical inputs to PT neurons from controls and 6-OHDA mice. Open and filled circles indicate individual events of asynchronous EPSCs. **H-I**) Box plots showing unaltered frequency (H) and amplitude (I) of Sr^2+^-EPSCs at sensory cortical inputs to M1 PT neurons between groups. *, *p* < 0.05, ns, not significant.

### Loss of DA alters the structure of thalamocortical synapses to M1 PT neurons

To assess how loss of DA alters thalamocortical innervation of M1 PT neurons, brain sections from control and 6-OHDA mice were processed for immunohistochemical detection of vesicular glutamate transporter 2 (vGluT2), a marker of thalamic axon boutons in M1. Stereological methods were used to quantify vGluT2-immunoreactive (vGluT2-ir) puncta in the layer V of M1. The density of vGluT2-ir puncta was significantly decreased in slices from 6-OHDA mice relative to those from controls (control = 20.4 [16.9–22.9] million puncta/mm^3^, n = 20 slices/3 mice; 6-OHDA = 11.2 [8.5–13.9] million puncta/mm^3^, n = 20 slices/3 mice; *p* < 0.0001, MWU; **Figure 3A-C**). These data are consisted with findings from MPTP-treated NHP (Villalba et al., 2021). It has been reported that reduced vGluTs levels per terminal leads to a smaller vesicle size (Wojcik et al., 2004). Considering the unaltered quantal size of Sr^2+^-EPSCs (**Figure 2A, C**), we posit that the reduced vGluT2 density indicates a loss of presynaptic thalamic terminals, rather than a decreased vGluT2 protein expression per thalamic axon terminal.

**Figure 3.**
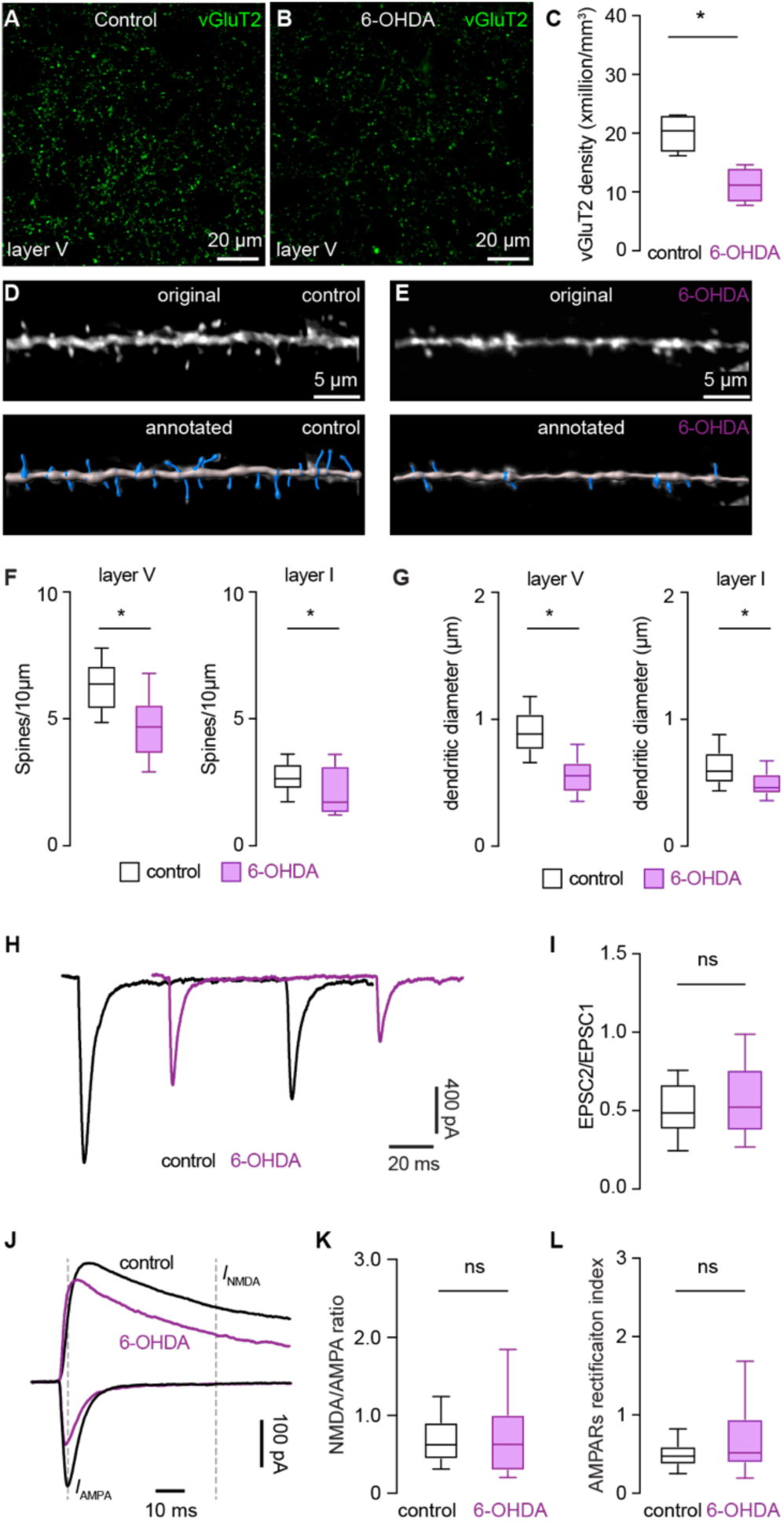
Degeneration of SNc DA neurons induces structural alterations of thalamocortical synapses. **A-C**) Representative confocal images of vGluT2-immunoreactivity in the layer 5 of M1 from controls (A) and 6-OHDA mice (B), and the summarized results showing a reduced vGluT2 density in the layer 5 of M1 following the SNc DA degeneration (**C**). **D-E**) Representative confocal images of segments of basal dendrites of M1 PT neurons from controls (D) and 6-OHDA mice (E). The bottom panels show the annotated dendrites using Imaris software. **F-G**) Summarized results showing decreased spine densities of L5 basal dendrites and L1 apical dendrites (F) and reduced dendritic diameters of L5 basal dendrites and L1 apical dendrites from 6-OHDA mice, relative to controls. **H-I**) Representative traces of EPSCs in response to paired pulses stimulation from controls and 6-OHDA mice (H) and the summarized results showing unaltered pared-pulse ratios between groups. **J-L**) Representative traces of optogenetically-evoked thalamic EPSCs recorded at -80 mV and +40 mV in PT neurons from controls and 6-OHDA mice. Dashed lines indicate where AMPARs- and NMDARs-mediated responses were measured to quantify NMDA/APMA ratio (K) and AMPARs rectification (L). Summarized results showing no change in the NMDA/AMPA ratio (K) and AMPARs rectification index (L) between groups.

Thalamic axon terminals target the dendritic spines and dendritic shaft of M1 PT neurons (Hooks et al., 2013; Kuramoto et al., 2009; Villalba et al., 2021). To interrogate postsynaptic structural changes associated with DA degeneration, the density of dendritic spines and the diameter of dendritic shaft of M1 PT neurons were assessed in slices from controls and 6-OHDA mice that received AAVrg-hSyn-eGFP injections into the pontine nuclei (**Figure 3D, E**). The density of spines on the basal dendrites in the layer V from 6-OHDA mice decreased significantly relative to those from controls (spine density of basal dendrites, control = 6.4 [5.4–7.1] per 10 μm, n = 53 segments/3 mice; 6-OHDA = 4.7 [3.6–5.5] per 10 μm, n = 47 segments/3 mice; *p* < 0.001, MWU; **Figure 3D-F**). Similarly, the densities of eGFP-labeled spines on the L1 apical dendrites also decreased significantly in slices from 6-OHDA mice relative to those from controls (spine density of L1 apical dendrites, controls = 2.6 [2.3–3.2] per 10 μm, n = 24 segments/3 mice; 6-OHDA = 1.7 [1.3–3.1] per 10 μm, n = 25 segments/3 mice; *p* = 0.02, MWU, **Figure 3F**). Further, the diameters of both L5 basal dendrites (control = 0.9 [0.77–1.06] μm, n = 262 samples/3 mice; 6-OHDA = 0.57 [0.44–0.67], n = 267 samples/3 mice; *p* < 0.0001, MWU; **Figure 3G**) and L1 apical dendrites (control = 0.59 [0.5–0.74] μm, n = 240 samples/3 mice; 6-OHDA = 0.46 [0.42–0.57], n = 240 samples/3 mice; *p* < 0.0001; MWU; **Figure 3G**) were decreased significantly in slices from 6-OHDA mice relative to those from controls. The above data suggest that SNc DA degeneration induces loss of dendritic spines and shrunk dendritic shafts of M1 PT neurons. Combining structural changes at both the pre- and post-synaptic sides (**Figure 3A-G**), we conclude that degeneration of SNc DA neurons induces a loss of thalamocortical synapses to M1 PT neurons.

### SNc DA degeneration does not alter postsynaptic receptor properties of thalamocortical synapses to M1 PT neurons

Next, we assessed function adaptations at thalamocortical inputs to M1 PT neurons following the loss of SNc DA neurons. There was no significant change in the paired pulse ratio (PPR) of thalamocortical oEPSCs in PT neurons from control and 6-OHDA mice (control = 0.49 [0.39–0.66], n= 62 neurons/7 mice; 6-OHDA = 0.52 [0.38–0.75], n = 73 neurons/8 mice; *p* = 0.1, MWU; **Figure 3H, I**). It suggests that the decreased thalamocortical projection to PT neurons was not due to a lower initial release probability at thalamic axon terminals.

To assess postsynaptic adaptations at thalamocortical synapses to PT neurons, we measured the ratio of EPSCs mediated by NMDA receptors (Rs) and AMPARs (NMDA/AMPA ratio) as well as AMPARs rectification properties between controls and 6-OHDA mice. There was no difference in the NMDA/AMPA ratio at thalamic projections to PT neurons between groups (control = 0.62 [0.44–0.91], n = 34 neurons/4 mice; 6-OHDA = 0.63 [0.29–1.0], n = 38 neurons/5 mice; *p* = 0.85, MWU; **Figure 3J, K**).

Moreover, there was no alteration in the inward rectification index of AMPARs at thalamic projections to PT neurons (control = 0.47 [0.37–0.6], n = 34 neurons/4 mice; 6-OHDA = 0.52 [0.39–0.94], n = 38 neurons/5 mice; *p* = 0.22, MWU; **Figure 3J, L**). These data suggest that SNc DA degeneration does not alter the relative contributions of postsynaptic AMPARs- and NMDARs-mediated transmission or change the subunit composition of AMPARs at thalamic projections to M1 PT neurons.

### SNc DA degeneration decreases the effectiveness of thalamic driving of M1 PT neuronal firing

Loss of SNc DA neurons selectively decreases the intrinsic excitability of M1 PT neurons (Chen et al., 2021). Together with the decreased thalamocortical synaptic strength of PT neurons, a corollary prediction would be a reduced effectiveness of thalamic driving of M1 PT neuronal firing following the loss of SNc DA neurons. To test this hypothesis, we recorded retrogradely labelled PT and IT neurons under current clamp mode and compared the effectiveness of thalamic excitation in driving the action potential (AP) firing in controls and 6-OHDA mice (**Figure 4A, D**). A 20-Hz optogenetic stimulation (1 ms duration) was delivered to activate ChR2-tagged thalamic axons in M1, mimicking the synchronized β band bursting pattern of thalamocortical activity observed in parkinsonian state (Brazhnik et al., 2016; Oswal et al., 2013). As predicted, the effectiveness of thalamic driving of PT neuronal firing decreased significantly in slices from 6-OHDA mice relative to those from controls (**Figure 4A-C**), as reflected by a decreased number of APs per stimulation (*p* < 0.0001 for group difference, two-way ANOVA followed by Sidak’s tests; control = 29 neurons/3 mice; 6-OHDA = 33 neurons/4 mice; **Figure 4A, B**) and a decreased AP firing probability (*p* < 0.0001 for group difference, two-way ANOVA followed by Sidak’s tests; control = 29 neurons/3 mice; 6-OHDA = 33 neurons/4 mice; **Figure 4A, C**). In contrast, we did not detect alterations in the effectiveness of thalamic driving of IT neurons firing between groups (*p* = 0.56 for group difference on AP probability; *p* = 0.36 for group difference on the number of APs; control = 19 neurons/3 mice; 6-OHDA = 17 neurons/3 mice; two-way ANOVA, **Figure 4D-F**). Thus, these data suggest that loss of DA selectively disrupts the effectiveness of thalamic excitation to drive PT neurons firing in M1.

**Figure 4.**
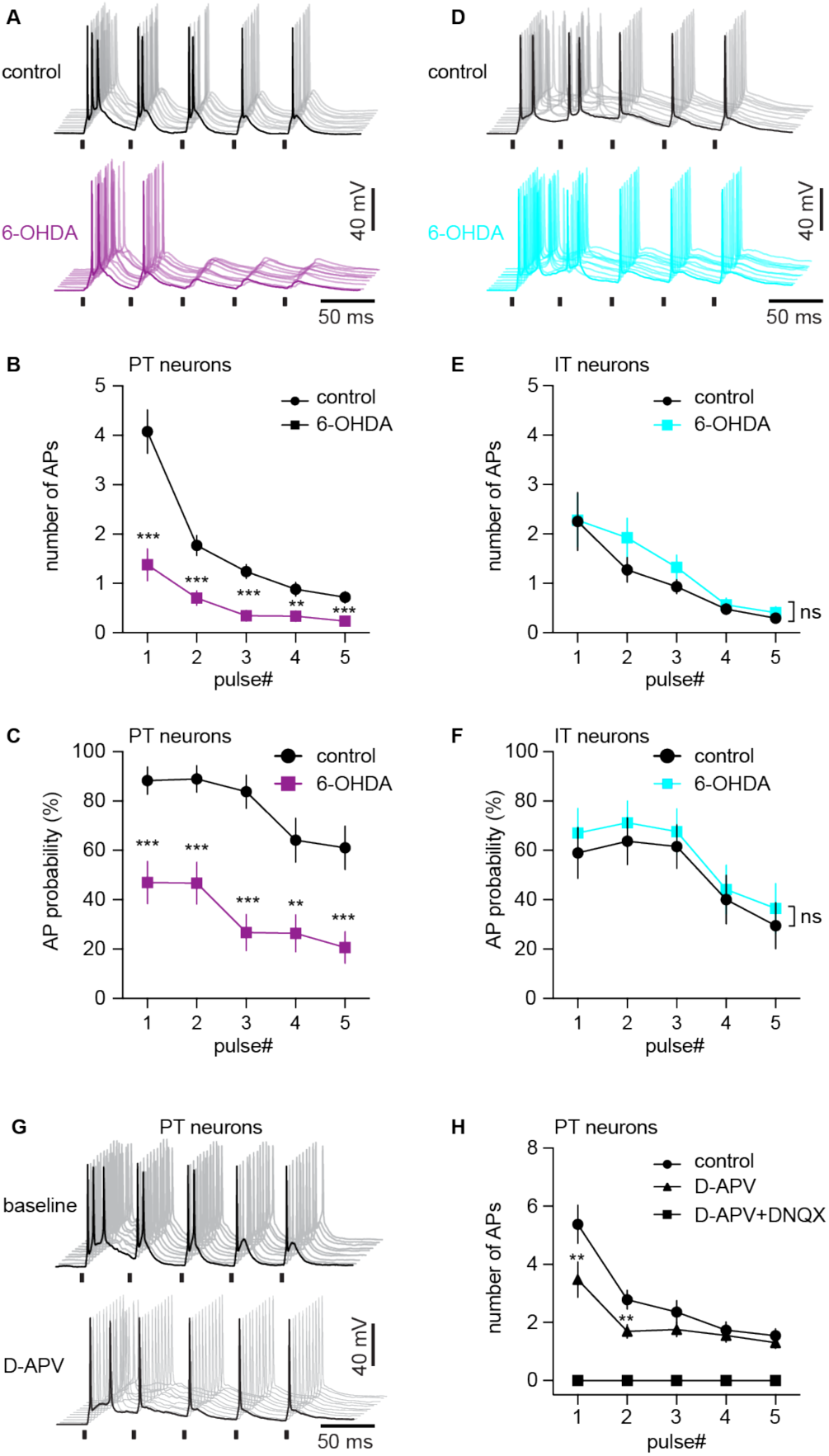
SNc DA degeneration decreases the effectiveness of thalamic driving of M1 PT neuronal firing. **A**) Representative traces of AP firing of M1 PT neurons from controls and 6-OHDA mice in response to 20 Hz optogenetic stimulation. AP traces from 10 trials were aligned for both groups. **B-C**) Summarized results show that both the number (B) and probability (C) of synaptically-generated APs in M1 PT neurons decreased significantly in 6-OHDA mice relative to controls. **D**) Representative traces of AP firing of M1 IT neurons from controls and 6-OHDA mice in response to 20 Hz optogenetic stimulation. **E-F**) Summarized results show that the number (E) and probability (F) of synaptically-generated APs in M1 IT neurons was not altered in 6-OHDA mice relative to controls. **G**) Representative traces of AP firing of M1 PT neurons in response to 20 Hz optogenetic stimulation prior to and after D-APV application. **H**) Summarized results show that NMDARs blockade using D-APV decreased the number of APs of PT neurons generated by thalamic stimulation.

To determine the relative contributions of AMPARs- and NMDARs-mediated responses to thalamic driving of PT neurons firings, pharmacological approach was used to assess the impact of NMDARs and AMPARs blockades on the number of synaptically generated APs from PT neurons in control mice. The number of APs generated by PT neurons in response to optogenetic stimulation of thalamic axon terminals was significantly reduced by blockade of NMDARs using D-APV (50 μM), particularly the number of APs generated by the first two stimulations (two-way ANOVA followed by Sidak’s tests; 13 neurons/3 mice **Figure 4G**). Synaptically generated APs of PT neurons were abolished by the subsequent blockade of both NMDARs and AMPARs using a mixture of D-APV (50 μΜ) and DNQX (20 μM) (**Figure 4H**). These data suggest that both the NMDARs and AMPARs mediate thalamic driving of M1 PT neuron firing in control mice. Altogether, the above results suggest that SNc DA degeneration selectively decreases thalamocortical connectivity to M1 PT neurons and impairs the effectiveness of thalamic driving of PT neuronal firing.

### Distinct properties of postsynaptic receptors at thalamic and sensory cortical inputs to M1 PT neurons

NMDA receptors play a critical role in thalamocortical synaptic integration and plasticity during learning motor skills and contribute to adaptative changes in neurological disorders (Guo et al., 2015b; Lafourcade et al., 2022; Miller et al., 2017). Next, we tested the hypothesis that NMDARs at thalamocortical synapses mediate the selective downregulation of thalamic excitation of PT neurons following the loss of SNc DA neurons. To do this, we first measured the NMDA/AMPA ratio at thalamic projections to PT and IT neurons (i.e., VM-PT and VM-IT synapses, respectively) as well as sensory cortical synapses to PT neurons (i.e., SC-PT synapse) in controls. Interestingly, we found a significantly higher NMDA/AMPA ratio at the thalamic synapses to PT neurons, relative to thalamic projection to IT neurons or SC-PT synapses (NMDA/AMPA ratio, VM-PT synapses = 0.62 [0.44–0.91], n = 34 neurons/4 mice; VM-IT synapse = 0.22 [0.16–0.33], n = 25 neurons/3 mice; SC-PT synapses = 0.34 [0.22–0.55], n = 19 neurons/3 mice; *p* < 0.0001; Kruskal-Wallis test followed by Dunn’s test, **Figure 5A, B**). These data suggest that NMDARs are functionally enriched at VM-PT synapses relative to SC-PT or VM-IT synapses. In addition, there was a stronger inward rectification of AMPARs at SC-PT synapses relative to thalamic projections to PT or IT neurons (AMPARs rectification index, VM-PT synapses = 0.47 [0.37–0.60], n = 34 neurons/4 mice; VM-IT synapse = 0.91 [0.54–1.4], n = 25 neurons/3 mice; SC-PT synapses = 1.23 [0.97–2.32], n = 19 neurons/3 mice; *p* < 0.0001; Kruskal-Wallis test followed by Dunn’s test; **Figure 5A, C**). These data suggest GluA2-lacking, Ca^2+^-permeable AMPARs are enriched at SC-PT synapses relative to those from the motor thalamus to M1 PT or IT neurons in control mice.

**Figure 5.**
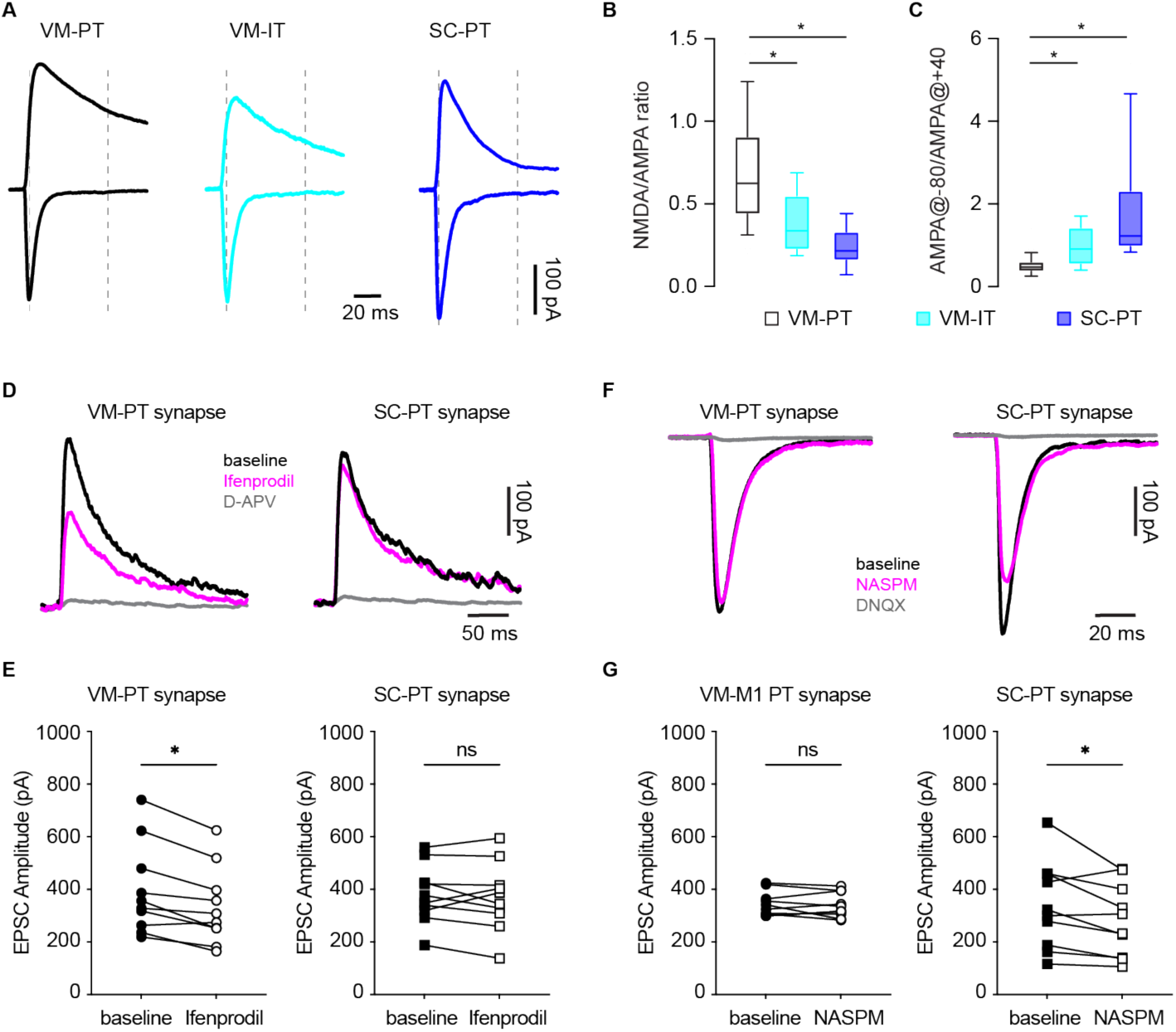
Distinct postsynaptic receptor properties at thalamic and sensory cortical inputs to PT neurons. **A**) Representative traces of AMPARs- and NMDARs-mediated responses from different synapses in M1. **B-C**) Box plots showing difference in the NMDA/AMPA ratios (**B**) and AMPARs rectification properties (**C**) of different synapses in M1. **D-E**) Pharmacology studies showing different sensitivities of NMDARs-mediated transmission at thalamic and sensory cortical synapses to ifenprodil. **D**) Representative traces of isolated NMDARs-mediated EPSCs from VM-PT and SC-PT synapses at baseline, in ifenprodil, and in ifenprodil plus D-APV. **E**) Box plots showing the effects of ifenprodil to the amplitude of NMDARs-mediated EPSCs at VM-PT (left) and SC-PT (right) synapses. **F**) Representative traces of AMPARs-mediated EPSCs from VM-PT and SC-PT synapses at baseline, in NASPM, and in NASPM plus DNQX. **G**) Box plots showing the effects of NASPM to the amplitude of AMPARs-mediated EPSCs at VM-PT (left) and SC-PT (right) synapses.

Next, pharmacological approaches were used to further confirm the synapse-specific properties of different inputs to M1 PT neurons in control mice. A large body of evidence suggests that GluN2B subunit-containing NMDA receptors with larger Ca^2+^ conductance enriched at thalamocortical synapses in the prefrontal cortex and play a critical role in pathological processes in neurological diseases (Miller et al., 2017; Wang et al., 2008). Consistently, a noncompetitive antagonist of GluN2B-containing NMDARs ifenprodil (3 μM) significantly and consistently reduced the amplitude of NMDARs-EPSCs at VM-PT synapses (baseline = 342 [256–516] pA, ifenprodil = 292 [234–428] pA; n = 10 neurons/4 mice; *p* = 0.004, WSR; **Figure 5D, E**), but not did not produce consistent effect to the NMDA-EPSCs amplitude of SC-PT synapses (baseline = 363 [313–452] pA, ifenprodil = 366 [296–443] pA; n = 10 neurons/3 mice; *p* = 0.6, WSR; **Figure 5D, E**). In addition, to quantitively assess the functional contribution of GluA2-lacking, Ca^2+^-permeable AMPARs at different inputs, a selective antagonist1-Naphthyl acetyl spermine trihydrochloride (NASPM) was used. Bath application of NASPM (100 μΜ) consistently decreased the amplitude of AMPARs-EPSCs of SC-PT synapses (baseline = 311 [181–459] pA; NASPM = 269 [106–419] pA; n = 10 neurons/ 4 mice; *p* = 0.02, WSR; **Figure 5F, G**), but did not produce significant effect to the amplitude of VM-PT EPSCs (baseline = 333 [305–378] pA; NASPM = 327 [301–395] pA; n = 10 neurons/ 4 mice; *p* = 0.6, WSR; **Figure 5F, G**).

Together, we conclude that GluN2B-containing NMDARs are enriched at thalamic transmission to M1 PT neurons, and that GluA2-lacking, Ca^2+^-permeable AMPARs functionally contribute more to sensory cortical transmission to M1 PT neurons under normal state.

### NMDARs mediate the decreased thalamocortical transmission to M1 PT neurons

Prolonged activation of NMDARs by exogenously application of NMDA is sufficient to mimic NMDAR-dependent synaptic and cellular plasticity in basal ganglia nuclei (Chu et al., 2017; McIver et al., 2019). Given the enrichment of NMDARs at VM-PT synapses, we further assessed if NMDARs activation mediates the decreased thalamocortical synaptic strength to M1 PT neurons in parkinsonian state. Thus, brain slices from controls and 6-OHDA mice were incubated with (1) normal ACSF, (2) NMDA-containing ACSF (1 hour at 25 μM), or (3) D-APV/NMDA-containing ACSF for hours (Chu et al., 2017), prior to physiological and anatomical studies (**Figure 6A**). In brain slices from vehicle-injected control mice, NMDA incubation *ex vivo* reduced the amplitude of thalamic oEPSCs of M1 PT neurons (*p* < 0.0001, two-way ANOVA followed by Sidak’s tests; **Figure 6B, D**), and the effects of NMDA could be abolished by D-APV and NMDA co-incubation (**Supplementary Figure 1**). Interestingly, *ex vivo* NMDA incubation had no effect on the amplitude of thalamic oEPSCs of M1 PT neurons from 6-OHDA mice (*p* > 0.79; **Figure 6C, E**). These data suggest that chronic NMDARs activation is sufficient to decrease the thalamic excitation to M1 PT neurons from control mice, but such effects are occluded by 6-OHDA-induecd DA degeneration *in vivo*.

**Figure 6.**
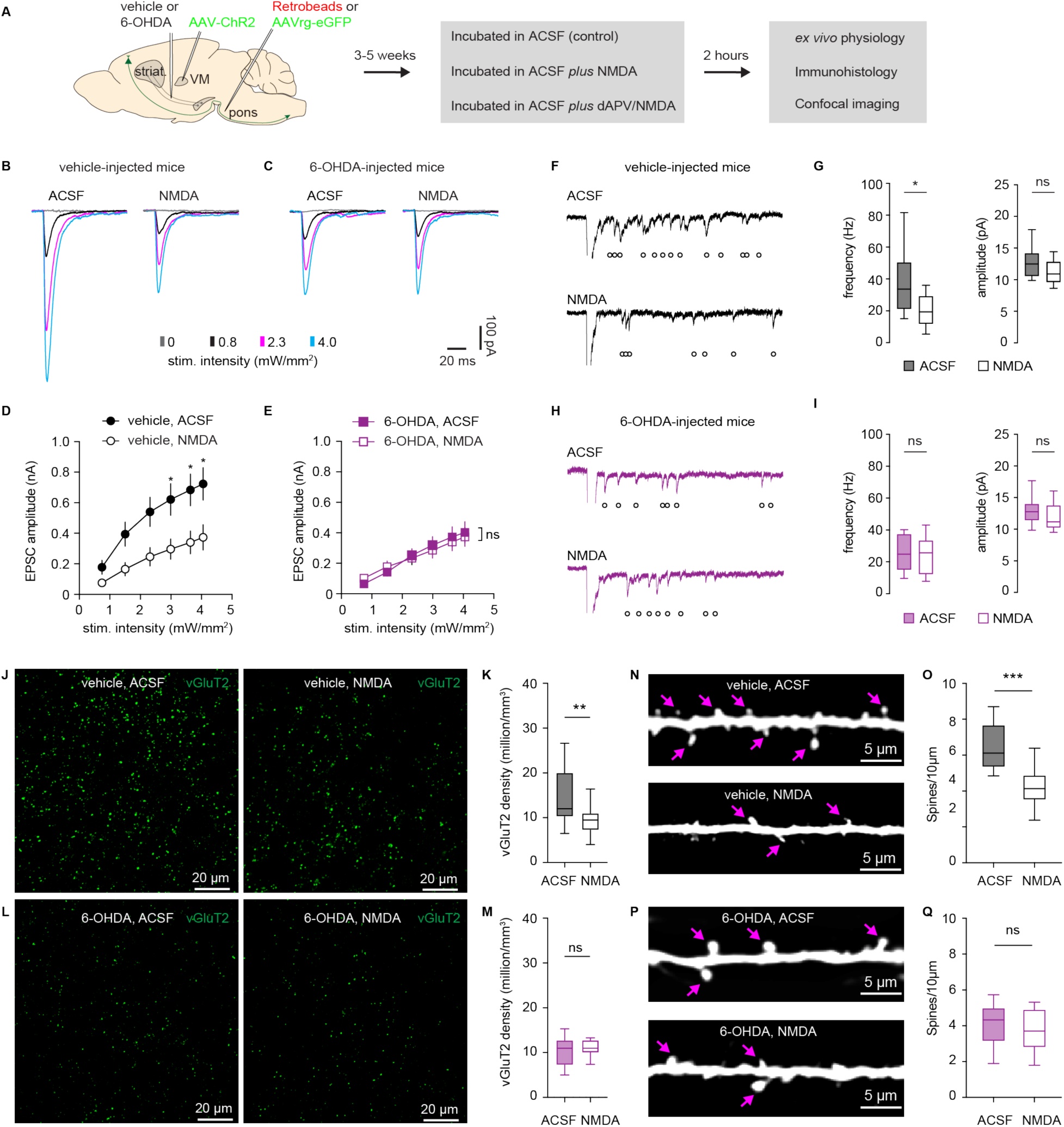
Excessive NMDARs activation mediates the impaired thalamocortical transmission following SNc DA degeneration. **A**) Experimental design. **B-C**) Representative traces of optogenetically-evoked thalamocortical EPSCs of PT neurons from vehicle-injected controls (B) and 6-OHDA-injected mice (C), which were either incubated in normal ACSF (control) or ACSF containing NMDA (25 μM). **D-E**) Summarized results show that NMDA incubation decreased the amplitude of oEPSCs from slices of vehicle-injected controls (D), but not those from slices of 6-OHDA-injected mice (E). **F-G**) NMDA incubation decreased the frequency but not the amplitude of optogenetically-evoked Sr^2+^-EPSCs in slice from vehicle-injected controls. **F**) Representative traces of Sr^2+^-EPSCs. **G**) Summarized results. **H-I**) NMDA incubation did not alter the frequency and the amplitude of optogenetically-evoked Sr^2+^-EPSCs in slice from 6-OHDA-injected mice. **H**) Representative traces of Sr^2+^-EPSCs. **I**) Summarized results. **J**) Representative confocal images of vGluT2 immunoreactivity from ACSF- and NMDA-treated slices of controls. **K**) Box plot shows that NMDA incubation reduced vGluT2 densities in slices of controls. **L**) Representative confocal images of vGluT2 immunoreactivity from ACSF- and NMDA-treated slices of 6-OHDA. **M**) Box plot shows that NMDA incubation did not affect vGluT2 densities in slices of 6-OHDA mice. **N**) Representative confocal images of M1 L5 dendritic spines from ACSF- and NMDA-treated slices of controls. **O**) Box plot shows that NMDA incubation reduced L5 spine densities in slices of controls. **P**) Representative confocal images of M1 L5 dendritic spines from ACSF- and NMDA-treated slices of 6-OHDA mice. **Q**) Box plot shows that NMDA incubation did not alter L5 spine densities in slices from 6-OHDA mice. Arrows in (N and P) indicate representative spines on dendrites.

Since the degeneration of SNc DA neurons decreases the number of functional thalamic inputs to M1 PT neurons (**Figures 2 & 3**), we further studied the impact of exogenous NMDA incubation on Sr^2+^-induced quantal glutamate release at thalamocortical synapses to PT neurons. In slices from vehicle-injected controls, NMDA incubation significantly decreased the frequency, but not the amplitude of optogenetically evoked thalamic Sr^2+^-EPSCs (frequency of Sr^2+^-EPSCs in vehicle-injected mice, control = 33.6 [21.1–50.4] Hz, n = 24 neurons/3 mice; NMDA = 10.9 [9.6–12.9] pA; n = 24 neurons/3 mice; *p* = 0.001, MWU; amplitude of Sr^2+^-EPSCs in vehicle-injected mice, control = 12.5 [10.5–14.2] pA, n = 24 neurons/3 mice; NMDA = 19.4 [11.8–29.2] Hz, n = 24 neurons/3 mice; *p* = 0.1, MWU; **Figure 6F, G**). In contrast, in slices from 6-OHDA mice, NMDA incubation did not alter the frequency or the amplitude of optogenetically-evoked thalamic Sr^2+^-EPSCs of PT neurons (frequency of Sr^2+^-EPSCs in 6-OHDA mice, control = 24.8 [15–37.4] Hz, n = 17 neurons/3 mice; NMDA = 25.6 [12.2–33.4] pA, n = 17 neurons/3 mice; *p* = 0.6, MWU; amplitude of Sr^2+^-EPSCs in 6-OHDA mice; control = 12.9 [11.3–14.0] pA, n = 17 neurons/3 mice; NMDA = 11.2 [10.3–13.8] Hz, n = 27 neurons/3 mice; *p* = 0.21, MWU; **Figure 6H, I**). The above data suggest that NMDARs activation reduces the number of functional thalamocortical inputs to M1 PT neurons, but this effect was occluded in 6-OHDA mice.

Next, we studied whether *ex vivo* NMDA incubation could induce structural changes at thalamocortical synapses to PT neurons. A separate set of brain slices from controls and 6-OHDA mice were incubated with ACSF or NMDA-containing ACSF for 2 hours and then fixed with 4% PFA for immunohistochemistry studies. We found that *ex vivo* NMDA incubation robustly decreased M1 L5 vGluT2 densities from slices of controls, relative to those incubated in ACSF (vGluT2 densities from controls, ACSF = 12 [10–14.8] million/mm^3^, n = 20 samples/3 mice; NMDA = 9 [7–10.8] million/mm^3^, n = 16 samples/3 mice; *p* = 0.0016, MWU; **Figure 6J, K**). However, NMDA incubation did not alter M1 L5 vGluT2 densities in slices from 6-OHDA mice (vGluT2 densities from 6-OHDA mice, ACSF = 11 [8–12] million/mm^3^, 20 samples/3 mice; NMDA = 11 [9–12] million/mm^3^, 20 samples/3 mice; *p* = 0.7, MWU; **Figure 6L, M**). In addition, NMDA incubation also decreased the spine densities of L5 basal dendrites of retrogradely labeled PT neurons from controls (spine density from controls, ACSF = 6 [5.4–7.6] spines/10 μm, n = 24 samples/3 mice; and NMDA = 4.1 [3.5–4.9] spines/10 μm, n = 25 samples/3 mice; *p* < 0.0001, MWU; Figure N, O); but NMDA incubation did not alter spine densities of PT neurons in slices from 6-OHDA mice (spine density from 6-OHDA mice, ACSF = 4.3 [3.2–5] spines/10 μm, n = 29 samples/3 mice; NMDA = 3.7 [2.8–4.9] spines/10 μm, n = 31 samples/3 mice; *p* = 0.4, MWU; **Figure 6P, Q**). Altogether, our physiological and anatomical results suggest that excessive NMDARs activation is sufficient to induce a reduced thalamocortical connection to M1 PT neurons, and that DA degeneration likely recruits the NMDA-mediated mechanisms, leading to the decreased thalamic excitation to M1 PT neurons in 6-OHDA mice.

Our recent work also showed that loss of DA decreases the intrinsic excitability of M1 PT neurons (Chen et al., 2021). Whether the intrinsic and synaptic adaptations in PT neurons share the same NMDA-mediated mechanisms remain unknown. To address this question, we studied the impact of NMDA incubation on cellular excitability of M1 PT neurons from control mice. Interestingly, exogenous NMDA incubation did not alter the cellular excitability of PT neurons in slices from controls (**Supplementary Figure 2**). The above data indicate chronic NMDARs activation was not sufficient or necessary to trigger intrinsic adaptations in M1 PT neurons. Taken together, these data suggest that (1) NMDARs activation *ex vivo* was sufficient to decrease thalamocortical transmission in slice from control mice; (2) in DA depleted 6-OHDA mice NMDARs-mediated reduction of thalamocortical transmission to PT neurons *in vivo* occluded the effect of NMDAR activation *ex vivo*; and (3) distinct molecular mechanisms mediate the intrinsic and synaptic adaptations in M1 PT neurons following the degeneration of SNc DA neurons.

### Basal ganglia outputs drive the decreased thalamocortical transmission following the degeneration of SNc DA neurons

Cortico-basal ganglia network shows an exaggerated oscillatory and bursting pattern of activity following the loss of SNc DA neurons (Galvan and Wichmann, 2008; Oswal et al., 2013). We posited that the rhythmic and bursting outputs of the basal ganglia promote NMDARs activation at the thalamocortical synapses to M1 PT neurons, and that prolonged NMDARs activation further contributes to the decreased thalamocortical excitation to PT neurons. Thus, we employed chemogenetic approaches to suppress basal ganglia output *in vivo* and studied whether M1 circuitry dysfunction in 6-OHDA mice could be rescued by inhibiting pathologic basal ganglia outputs.

PV-Cre knockin mice were unilaterally injected with vehicle or 6-OHDA into the MFB, plus AAV9-hSyn-FLEX-hM4Di-mCherry into the SNr of the same hemisphere for chemogenetic manipulation of basal ganglia outputs (**Figure 7A**). Immunofluorescence staining showed a selective expression of hM4Di (Gi)-DREADDS on SNr PV-expressing cells (**Figure 7A**). There was no difference in the number of hM4Di-expressing cells in the SNr between 6-OHDA mice and controls (controls = 276 [160–357] neurons/slice, n = 20 slices/10 mice; 6-OHDA = 233 [170–271] neurons/slice, n = 23 slices/12 mice; *p* = 0.08, MWU). Loose-sealed cell-attached recordings showed that bath application of DREADDS agonist CNO (10 μΜ) effectively suppressed the frequency and regularity of autonomous firing of SNR neurons from 6-OHDA mice (frequency, baseline = 5.2 [4.3– 7.5] Hz, CNO = 3.9 [0.27–5.6] Hz; n = 10 neurons/3 mice; *p* = 0.02, WSR; **Figure 7C**; CV of ISIs, baseline = 0.27 [0.13–0.49], CNO = 0.61 [0.19–1.35]; n = 10 neurons/3 mice; *p* = 0.016, WSR; **Figure 7D**).

**Figure 7.**
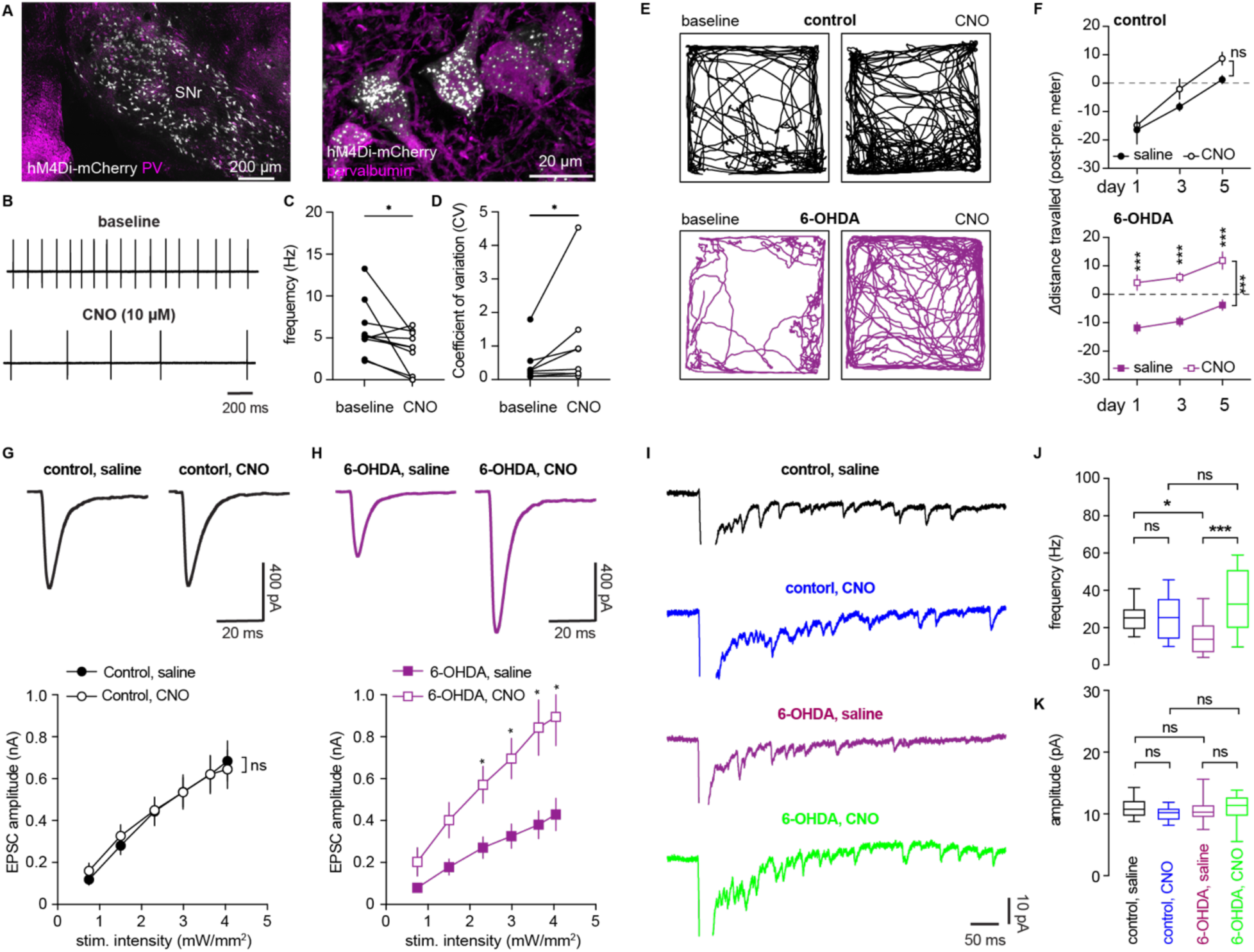
Figure 7. Basal ganglia output drives the synaptic adaptations in M1 following the loss of SNc DA neurons. **A)** Representative confocal images showing hM4Di-mcherry expression in the PV-expressing SNr neurons under low and high magnifications. **B)** Representative traces of the autonomous firing of SNr neurons from a 6-OHDA mice prior to and after CNO application. **C-D)** Box plots showing that hM4Di activation by CNO effectively suppressed the frequency (C) and regularity (D) of SNr neurons firing. **E)** Representative track plots showing the effects of CNO injections to locomotor activity of controls and 6- OHDA mice in an open field on day 5. **F)** Summarized results showing the difference of distance travelled in the open field between pre- and post-saline/CNO injections in controls and 6-OHDA mice. Behavioral tested were conducted on days 1, 3 and 5. **G-H)** hM4Di activation by CNO injection did not alter the amplitude of optogenetically thalamocortical EPSCs in PT neurons from controls (G), but it induced an increased optogenetically thalamocortical EPSCs in PT neurons from 6-OHDA mice (H). Top, representative traces of oEPSCs in PT neurons; Bottom, summarized results. **I)** Representative traces of optogenetically-evoked Sr^2+^-EPSCs in controls and 6-OHDA mice treated with saline or CNO. **J)** Summarized results showing CNO injection did not alter the frequency of Sr^2+^- EPSCs in slices from controls, but decreased the frequency of Sr^2+^-EPSCs in slices from 6-OHDA mice. **K)** Summarized results showing CNO injection did not alter the amplitude of Sr^2+^-EPSCs in slices from either controls or 6-OHDA mice.

PV-Cre mice with SNr hM4Di expression were injected with either CNO (1.0 mg/kg, S.C., at ∼12 hrs interval between injections) or saline for 5 consecutive days to continuously suppress abnormal basal ganglia outputs, or control for subcutaneous injections. Open filed locomotor activities were measured on days 1, 3 and 5. CNO injection tended to increase the distance travelled of control mice in an open field (**Figure 7E**), but such effects were not distinguishable from saline-injected control mice (**Figure 7F**). This is perhaps because that CNO’s effects to neuronal activity of the unilateral SNr of DA intact mice were not sufficient to robustly affect general motor activity. In contrast, CNO injection significantly increased the locomotor activity of 6-OHDA mice relative to saline injected mice (**Figure 7E**, **F**). These data indicate that the SNr neurons of 6-OHDA mice were more sensitive to hM4Di activation and that chemogenetic suppression of SNr output of 6-OHDA mice was associated with immediate behavioral effects. These results also confirm that chemogenetic approach could effectively suppress basal ganglia outputs and led to physiological and behavioral phenotypes.

To determine whether suppression of basal ganglia outputs could rescue adaptive changes of M1 circuits, the same cohort of mice described above also received retrobeads injection into the pons to retrogradely label M1 PT neurons and ChR2(H134R)-eYFP injection into the motor thalamus for optogenetic stimulation of thalamocortical synapses to PT neurons. In vehicle-injected control mice, we found that chemogenetic suppression of SNr activity through repetitive CNO injection did not alter the amplitude of optogenetically evoked thalamic oEPSCs in PT neuros (*p* = 0.9, two-way ANOVA; **Figure 7G**). In contrast, repetitive CNO injection significantly increased the amplitude of optogenetically evoked thalamic oEPSCs in PT neurons of 6-OHDA mice (*p* = 0.007, two-way ANOVA followed by Sidak’s; **Figure 7H**).

Further, we assessed the effects of chemogenetic suppression of SNr activity to quantal properties of thalamocortical transmission to M1 PT neurons by replacing extracellular Ca^2+^ with Sr^2+^. Chemogenetic suppression of SNr activity through repetitive CNO injection did not alter either the frequency or the amplitude of optogenetically-evoked thalamic Sr^2+^-EPSCs in PT neuros of vehicle-injected control mice (frequency of Sr^2+^-EPSCs, control/saline = 25 [19–30] Hz, control/CNO = 25 [14–35] Hz; *p* > 0.9; amplitude of Sr^2+^-EPSCs, control/saline = 10.7 [9.7–12.1] pA, n= 18 neurons/3 mice; control/CNO = 10.2 [9.1–10.9] pA, n = 18 neurons/3 mice; *p* = 0.8, Kruskal-Wallis test followed by Dunn’s tests, **Figure 7I-K**). In contrast, repetitive CNO injection significantly increased the frequency, but not the amplitude, of optogenetically evoked thalamic Sr^2+^-EPSCs in PT neurons of 6-OHDA mice (frequency of Sr^2+^-EPSCs, 6-OHDA/saline = 13.8 [6.7–21.3] Hz, 6-OHDA/CNO = 32.4 [19.6–50.8] Hz, *p* = 0.0001; amplitude of Sr^2+^-EPSCs, 6-OHDA/saline = 10.3 [9.5–11.4] pA, n= 22 neurons/3 mice; 6-OHDA/CNO = 11.4 [9.7–12.7] pA, n = 19 neurons/3 mice; *p* = 0.95, Kruskal-Wallis test followed by Dunn’s tests, **Figure 7I-K**). Together, these data suggest that chemogenetic suppression of basal ganglia outputs could rescue the decreased thalamocortical transmission to M1 PT neurons in 6-OHDA mice.

Our recent study reported a decreased intrinsic excitability of M1 PT neurons in parkinsonism (Chen et al., 2021). To interrogate the impact of chemogenetic suppression of basal ganglia output on cellular excitability of M1 PT neurons, retrogradely labelled PT neurons were targeted for current clamp recording. In vehicle-injected control mice, continuous CNO injections for 5 days did not alter the intrinsic excitability of M1 PT neurons (*p* = 0.29, two-way ANOVA, **Supplementary Figure 3**). In 6-OHDA mice, CNO injections for 5 days tended to increase the excitability of M1 PT neurons, but the effects did not reach statistical significance (*p* = 0.18, two-way ANOVA, **Supplementary Figure 3**). These data suggesting that adaptation of the intrinsic excitability of PT neurons perhaps involves additional mechanisms.

## Discussion

The present study provides novel insights into motor cortical circuitry adaptations following the degeneration of SNc DA neurons in parkinsonism. We demonstrated that degeneration of SNc DA neurons induces cell-subtype- and input-specific downregulation of thalamic excitation to M1 PT neurons (**Figure 1**). Our physiological and anatomical analyses showed that the weakened thalamocortical synaptic strength is likely caused by a loss of thalamocortical synapses (**Figure 2–4**). We further found that NMDARs are enriched at thalamocortical synapses to M1 PT neurons (**Figure 5**). Prolonged NMDARs activation *ex vivo* was sufficient to decrease thalamocortical excitation to M1 PT neurons from control animals, but such effects could be abolished by D-APV pretreatment and were occluded in slices from 6-OHDA mice (**Figure 6**).

These data suggest that excessive NMDARs activation *in vivo* is involved in mediating the decreased thalamocortical connection strength following the degeneration of SNc DA neurons. In addition, we showed that chemogenetic suppression of the SNr activity can reverse the disrupted thalamocortical transmission in 6-OHDA mice (**Figure 7**), suggesting that pathological activity of basal ganglia perhaps drives cortical circuitry dysfunction following the degeneration of SNc DA neurons. Last, we showed that synaptic and intrinsic adaptations of M1 PT neutrons in parkinsonism do not share the common NMDARs-mediated molecular mechanisms.

### Motor cortical circuits dysfunction in parkinsonism

It has been long thought that the hypoactivity of the motor cortex in PD is due to direct and acute effects of an elevated basal ganglia inhibition following the degeneration of SNc DA neurons. However, this model can’t explain the cell-subtype-specific decrease of PT (but not IT) neuronal activity in DA-depleted animals (Pasquereau and Turner, 2011), particularly considering that PT and IT neurons in M1 receive similar levels of thalamic excitation (Hooks et al., 2013). In addition, M1 circuits faces several perturbations in parkinsonism, e.g., decreased dopaminergic/noradrenergic neuromodulation (Gaspar et al., 1991; Vitrac and Benoit-Marand, 2017) and elevated/pathologic basal ganglia outputs. Thus, it is plausible that such circuitry perturbations engage homeostatic plasticity mechanisms to stabilize the neuronal and circuit activity (Turrigiano, 2011). As expected, we observed a series of intrinsic (Chen et al., 2021) and synaptic adaptations in M1 following the degeneration of SNc DA neurons (**Figures 1–4**), which mainly occur in PT neurons. Thalamocortical transmission and plasticity play critical roles in the acquisition (Biane et al., 2016) and execution (Sauerbrei et al., 2020) of skilled motor activity. Thus, the decreased thalamocortical connection strength (**Figure 1**) could contribute to the reduced magnitude and disputed temporal pattern of PT neuronal activation *in vivo* at the onset of skilled movements in parkinsonian mice and NHP (Aeed et al., 2021; Hyland et al., 2019; Pasquereau et al., 2016).

The reduced frequency of PT neurons firing can be caused by combined effects of intrinsic and synaptic adaptations, but these adaptive changes still can’t provide meaningful explanation for the enhanced synchronized bursting pattern of activity of M1 PT neurons in parkinsonian animals (Goldberg et al., 2002; Pasquereau et al., 2016).

Future detailed circuitry analyses at cellular and synaptic levels using optogenetics-assisted circuitry interrogations and computational approaches are needed to provide a better understanding of motor cortical dysfunction in parkinsonism.

### Molecular and circuitry mechanisms underlying motor cortical dysfunction

Our data show that NMDARs are enriched at thalamocortical synapses M1 PT neurons (**Figure 5**). Such NMDARs enrichment can play critical role in the synaptic plasticity at thalamocortical synapses during motor learning (Biane et al., 2016; Bruno and Sakmann, 2006; Guo et al., 2015b; Xu et al., 2009). Based on these basic biophysical properties, we predicted that excessive activation of NMDARs during the synchronized bursting pattern of PT neuronal activity in parkinsonian state would mediate the pathological loss of thalamocortical connection in 6-OHDA mice. Consistently we showed that prolonged NMDARs activation is sufficient to decreased thalamocortical connections in controls, but not in the 6-OHDA mice (**Figure 6**). Similar NMDARs-mediated maladaptive changes have been reported in the subthalamic nucleus and the medial prefrontal cortex in neurological disorders (Chu et al., 2017; Miller et al., 2017). Therefore, compartment specific innervation of M1 PT neurons by the thalamic inputs and the unique postsynaptic receptor compositions can be key molecular and synaptic mechanisms mediating their selective downregulation in parkinsonian state (Lafourcade et al., 2022).

Chronic suppression of basal ganglia outputs in 6-OHDA mice rescued the decreased thalamocortical transmission, arguing that the excessive pathological signals from the basal ganglia drive the adaptive changes in the M1. Consistently, recent studies have reported that synchronized bursting pattern of basal ganglia outputs can induce rebound firing of downstream targets, e.g., neurons in the motor thalamus and superior colliculus, and trigger the associated motor activities (Kim et al., 2017; Villalobos and Basso, 2022). Thus, we posit that the excessive basal ganglia outputs synchronize motor thalamic neuronal activity *in vivo* (Brazhnik et al., 2016; Kim et al., 2017), leading to a prolonged activation of NMDARs at thalamocortical synapses in M1 and a subsequent downregulation of their functional connection.

## Conclusion

Altogether, we demonstrated that degeneration of SNc DA neurons in parkinsonism triggers cell-subtype- and synapse-specific adaptations in M1, which can be important in the pathophysiology of motor deficits in PD. Moreover, our studies also highlight that synaptic and intrinsic alterations in M1 are mediated by different circuitry and molecular mechanisms. While thalamocortical synaptic adaptations are likely caused by excessively synchronized basal ganglia outputs and the associated NMDARs activation, additional mechanisms seem to be involved into the intrinsic adaptations of M1 PT neurons following the loss of DA neurons. Further studies remain needed for better understanding of motor cortical dysfunction in parkinsonism.

## Materials and Methods

### Animals

Wild-type (WT) C57BL/6J mice of both sexes (3–4 month-old, RRID:IMSR_JAX:000664) were obtained from the Van Andel Research Institute vivarium internal colony and used in the study. Homozygous PV-Cre knock-in mice were originally purchased from Jackson laboratories (stock# 017320RRID: IMSR_JAX:017320, Bar Harbor, ME) and maintained at a C57/BL6J background in Van Andel Research Institute vivarium. Mouse genotyping was conducted through the service of Transnetyx, Inc (Cordova, TN). Mice were housed up to four animals per cage under a 12/12 h light/dark cycle with access to food and water *ad libitum* in accordance with NIH guidelines for care and use of animals. All animal studies were reviewed and approved by the Institutional Animal Care and Use Committee at Van Andel Research Institute (reference#: 22-02-006).

### Stereotaxic surgery for 6-OHDA and virus injections

Mice were randomly assigned to different treatment groups. Mice were mounted and secured in a stereotaxic frame (Kopf) under 2% isoflurane anesthesia. Throughout the procedure, body temperature of the mouse was maintained by a thermostatic heating pad. Once the skull was opened, small holes were drilled above the pre-determined targets to inject: 1) 6-OHDA (3–4 μg) into the medial forebrain bundle (MFB, from bregma (in mm), anterior-posterior (AP) = -0.7, mediolateral (ML) = +1.2, and dorsoventral (DV) = -4.7 from the brain surface) to induce unilateral degeneration of nigrostriatal pathway, or vehicle into the MFB to serve as vehicle-injected controls; 2) retrobeads (200 nl, 10x dilution) into the ipsilateral pontine nuclei (from bregma (in mm), AP = -5.0, ML = +0.6, DV = -5.0) or contralateral striatum (from bregma (in mm), AP = +0.2, ML = -2.3, DV = -2.9) to retrogradely label PT and IT neurons, respectively; and 3) AAV9-hSyn-ChR2(H134R)-eYFP (RRID:Addgene_127090, 300 nl at a titer of 3.6×10^12^ GC/ml) into the ventral medial region of the thalamus (from bregma (in mm), AP = -1.6, ML = 0.8, DV = -4.3) to label thalamocortical axon terminals for optogenetics studies. A small cohort of mice also received pAAV9-hSyn-DIO-hM4D(Gi)-mCherry (RRID:Addgene_44362, 300 nl at a titer of 2.2×10^12^ GC/ml) injection into the substantia nigra pars reticulata (from bregma (in mm), AP = -3.4, ML = +1.4, DV = -4.6) for chemogenetic studies. For morphology studies, AAVrg-hSyn-eGFP (RRID:Addgene_50465, 300 nl at 2x 10^13^ GC/ml) were injected into the pontine nuclei to regrogradely label PT neuron in the motor cortex. All injection were performed using a 10 μl syringe (Hamilton, Reno, NV) mounted on a motorized microinjector (Stoelting, Wood Dale, IL) at a speed of 100 nl/min. Details of the stereotaxic injections procedure can be found on Protocol.io (dx.doi.org/10.17504/protocols.io.rm7vzye28lx1/v1)

### Slice preparation for electrophysiology

3–4 weeks post-injections, brain slices were prepared for electrophysiology. Mice were anesthetized with avertin (250–300 mg/kg), followed by transcardial perfusion with ice-cold sucrose-based artificial cerebrospinal fluid (aCSF) equilibrated with 95% O_2_/5% CO_2_ and containing (in mM): 230 sucrose, 26 NaHCO_3_, 10 glucose, 10 MgSO4, 2.5 KCl, 1.25 NaH_2_PO_4_, 0.5 CaCl_2_, 1 sodium pyruvate and 0.005 L-glutathione. Coronal brain sections (250 μm) containing the primary motor cortex were prepared using a vibratome (VT1200s, Leica, RRID:SCR_018453) in the same sucrose-based solution that was maintained at ∼4 °C using a recirculating chiller (FL300, Julabo, Allentown, PA). Slices were then held in aCSF equilibrated with 95% O_2_/5% CO_2_ and containing (in mM): 126 NaCl, 26 NaHCO_3_, 10 glucose, 2 MgSO_4_, 2.5 KCl, 1.25 NaH_2_PO_4_, 1 sodium pyruvate and 0.005 L-glutathione at 35 °C for 30 min for recovery and then at room temperature until electrophysiology recordings or NMDA incubation.

### Brain slice physiological recordings and optogenetics

Brain slices were placed in recording chamber with continuous perfusion of recording solution containing (in mM): 126 NaCl, 26 NaHCO_3_, 10 glucose, 3 KCl, 1.6 CaCl_2_, 1.5 MgSO_4_, and 1.25 NaH_2_PO_4_. The solution was equilibrated with 95% O_2_/5%CO_2_ and maintained at 33–34 °C using feedback controlled in-line heater (TC-324C, Warner Instruments). To assess synaptic strength, TTX (1 μΜ) and 4-AP (100 μM) were routinely included to isolate optogenetically evoked monosynaptic thalamocortical or corticocortical EPSCs in M1. To test the effectiveness of thalamic driving of M1 neuronal firing (Figure 4), TTX/4-AP were omitted from the recording solution and a selective GABA_A_ receptor antagonist SR-95531 (GABAzine) was added to prevent the recruitment of GABAergic inhibitions. Neurons were visualized using a CCD camera (SciCam Pro, Scientifica, UK) and SliceScope Pro 6000 system (Scientifica, UK) integrated with a BX51 upright microscope and motorized micromanipulators.

Retrogradely labelled neurons in the layer 5 were targeted under a 60x water immersion objective lens (Olympus, Japan) for whole-cell patch clamp recording using a MultiClamp 700B amplifier (RRID:SCR_018455) and Digidata 1550B controlled by pClamp 11 software (Molecular Devices, San Jose, CA; RRID:SCR_011323). Data were sampled at 50 KHz. Borosilicate glass pipettes (O.D. = 1.5 mm, I.D. = 0.86 mm, length = 10 cm, item#: BF150-86-10, Sutter Instruments, Novato, CA) were pulled by micropipette puller (P1000, Sutter instrument, Novato, CA; RRID:SCR_021042) and used for patch clamp recording with a resistance of 3–6 MOhm when filled with (1) cesium methanesulfonate-based internal solution of (in mM): 120 CH_3_O_3_SCs, 2.8 NaCl, 10 HEPES, 0.4 Na_4_-EGTA, 5 QX314-HBr, 5 phosphocreatine, 0.1 spermine, 4 ATP-Mg, and 0.4 GTP-Na (pH 7.3, 290 mOsm); or (2) K-gluconate based internal solution, containing (in mM): 140 K-gluconate, 3.8 NaCl, 1 MgCl_2_, 10 HEPES, 0.1 Na_4_-EGTA, 2 ATP-Mg, and 0.1 GTP-Na (pH 7.3, 290 mOsm). Optogenetic stimulation was delivered using a 478 nm LED (pE-300*^Ultra^*, CoolLED, UK; RRID:SCR_021972) through a 60x objective lens (Olympus). Electrophysiology data were analyzed and quantified using Clampfit 11.1 (Molecular Devices, San Jose, CA; RRID:SCR_011323).

### Immunohistochemistry

Brain tissues after electrophysiology brain slice preparation were fixed in 4% PFA in 0.1 M phosphate buffer overnight at 4 °C and then were resected into 70–100 μm slices using a VT1000s vibratome (Leica Biosystems, Deer Park, IL; RRID:SCR_016495) to prepare brain sections for immunohistochemistry of tyrosine hydroxylase (TH). In vGluT2 stereology and neuronal morphology studies, mice were perfused with ice-cold phosphate-buffered saline (PBS, pH = 7.4) for 5 min and subsequently with 4% PFA mice for 30 min. The brain was then extracted and saved in 4% PFA overnight at 4 °C before being resected into 70 μm slices using a VT1000s vibratome (Leica Biosystems, Deer Park, IL; RRID:SCR_016495) for immunohistochemistry or morphology studies.

Brain slices were rinsed 3x with PBS before being incubated with 0.2% Triton X-100 and 2% normal donkey serum (Sigma-Aldrich) for 60 min at room temperature. Brain slices were then incubated with primary antibodies, including mouse anti-TH (1:2000, cat# MAB318, Sigma-Aldrich; RRID:AB_2201528), or guinea pig anti-vGluT2 (cat# 135404, Synaptic Systems, RRID:AB_887884) for 48 hours at 4 °C or overnight at room temperature. Sections were then rinsed 3x with PBS and incubated with secondary antibodies for 90 min, including donkey anti-mouse Alexa Fluor 488 (Cat# 715-545-150; Jackson ImmunoResearch Labs, West Grove, PA; RRID:AB_2340846), donkey anti-mouse Alexa Fluor 594 (Cat# 715-585-150; Jackson ImmunoResearch Labs, West Grove, PA; RRID:AB_2340854), or donkey anti-guinea pig Alexa Fluor 488 (Cat# 706-545-148; Jackson ImmunoResearch Labs, West Grove, PA; RRID:AB_2340472), at room temperature before washing with PBS for 3 times. Brain sections were mounted with VECTASHIELD antifade mounting medium (Cat# H-1000, Vector Laboratories, Newark, CA; RRID:AB_2336789), were cover-slipped and sealed with nail polish. TH- and vGluT2-immunoreactivity (ir) was imaged using an Olympus DP80 camera through an Olympus BX63F microscope, or a confocal laser scanning microscope (A1R, Nikon, Japan; RRID:SCR_020317). TH-ir of control and 6-OHDA mice was quantified by normalizing the striatal TH-ir from the ipsilateral side to the value from the contralateral hemisphere, with a subtraction of background measured from the cerebral cortex.

### Animal behavior

To assess parkinsonian motor deficits, mice were subject to 1) open field locomotion test, where their locomotor activity and rotations were monitored for 10 min and quantified using Anymaze software (Stoelting, Wood Dale, IL); (2) cylinder test using a 600-ml glass beaker to assess the spontaneous forelimbs use during weight-bearing touches which were recorded using a digital camcorder at 60 fps (Panasonic, HC-V180K) and analyzed off-line by a researcher blinded to treatments. In chemogenetic studies, behavioral assessment was conducted 30 min after subcutaneous injection of saline-or water-soluble clozapine-n-oxide (CNO, 1 mg/kg body weight, cat# HB6149, HelloBio, Princeton, NJ).

### Confocal imaging and digital image analysis

vGluT2-ir was collected from the layers 5A and 5B under a 100x objective using a Nikon A1R confocal microscope. Z stack Images with a 0.15 μm interval were taken under the identical settings, including laser power, gain, pinhole size, and scanning speed etc. between animals. The density of vGluT2-ir was estimated based on the immunostaining signals between 5 and 8 μm below the surface of slices using the optical dissector method (West, 1999). For spine density analysis and dendritic examination, segments of basal dendrites were tracked down from the cell bodies of retrogradely labeled PT neurons in M1 and imaged under an oil-immersion 100x objective using the confocal microscope (NA = 1.45; x/y, 1024 X 1024 pixels; z step = 0.5 μm, A1R, Nikon). The distance from the soma to the basal dendrites analyzed was 50–100 μm. eGFP-labeled segments of the apical dendrites were selected from the layer I. To quantify spine density and dendritic diameter, only microscopic image data with ample eGFP expression and high signal-to-noise ratios were included. Spine density analysis was performed using Imaris software (version 9.3, Oxford, UK; RRID:SCR_007370). Briefly, 2-3 dendritic segments from the layer I and the layer V measuring 20–30 microns in length were reconstructed three-dimensionally using Imaris’ filament tracer function from each confocal image. Spines were then manually traced, reconstructed, and quantified. For accurate three-dimensionally reconstruction, both the dendrite segments and spines were recomputed automatically based on the manual traces. To quantify dendritic diameter, dendritic segments (30–40 μm in length) were traced and straightened using ImageJ (NIH, Bethesda, MD; RRID:SCR_003070). A grid (grid type: lines with 2 μm intervals; area per point: 5 μm^2^; center grid on image) was placed over top of the dendritic segments and cross-sectional measurements were taken along the dendrite at each grid mark (control: 240 and 262 measurements from 3 mice for the layer I and V, respectively; 6-OHDA: 240 and 267 measurements from 3 mice for the layer I and V, respectively).

### Statistics

Data were collected from 3-8 mice per study. Statistics analysis was performed in GraphPad Prism 9 (GraphPad Software, San Diego, CA; RRID:SCR_002798). Nonparametric Mann-Whitney U (MWU) test, Wilcox signed rank (WSR) test, and Kruskal–Wallis test followed by Dunn’s Multiple Comparisons (as indicated in the text) were used to compare the medians of 2 or 3 groups in order to minimize the assumption of dataset normality. Two-way ANOVA was used to compare effect of DA depletion or NMDA incubation on synaptic strength/AP firing across a range of stimulation intensities followed by Šídák’s multiple comparisons test. All statistic tests were two tailed with *p* values < 0.05 (*), 0.01 (**), or 0.001 (***) as thresholds for statistical significance. Data are reported as median plus interquartile ranges.

## Data availability

The datasets generated during and/or analyzed during the current study are available from the corresponding author on reasonable request.

## Author contribution

Investigation, formal analysis: L.C., S.D., R.D., and H.Y.C.; Conceptualization, funding acquisition, project administration, resources, supervision, visualization, writing – original draft: H.Y.C.; Writing – review& editing: L.C., S.D., and H.Y.C.

## Competing interests

The authors declare no competing interests.

## Acknowledgement

The authors thank the Van Andel Research Institute optical imaging core for the advanced confocal microscope, and Van Andel Research Institute vivarium for animal husbandry. The authors thank all members of the Chu lab at Van Andel Research Institute for thoughtful discussions. Funding: This work was supported by the National Institute of Neurological Disorders and Stroke (grant#: R01NS121371).

## Materials & Correspondence

Correspondence and material requests should be addressed to Dr. Hong-Yuan Chu

---

**Supplementary Figure 1.**
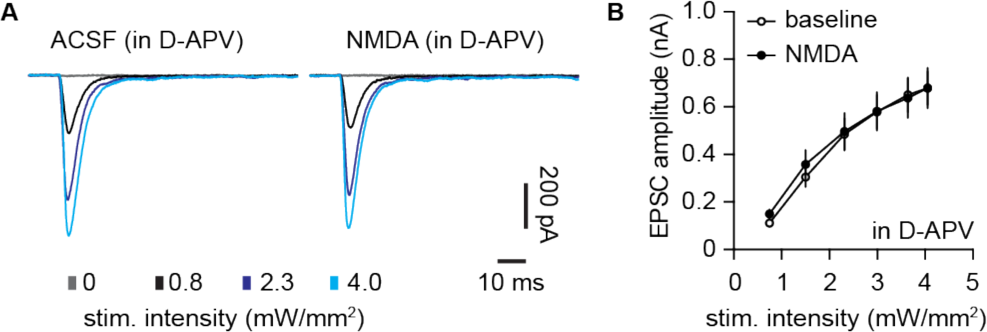
NMDA-induced reduction of thalamocortical transmission was blocked by D-APV. **A**) Representative traces of optogenetically-evoked thalamocortical EPSCs from PT neurons. **B**) Summarized data.

**Supplementary Figure 2.**
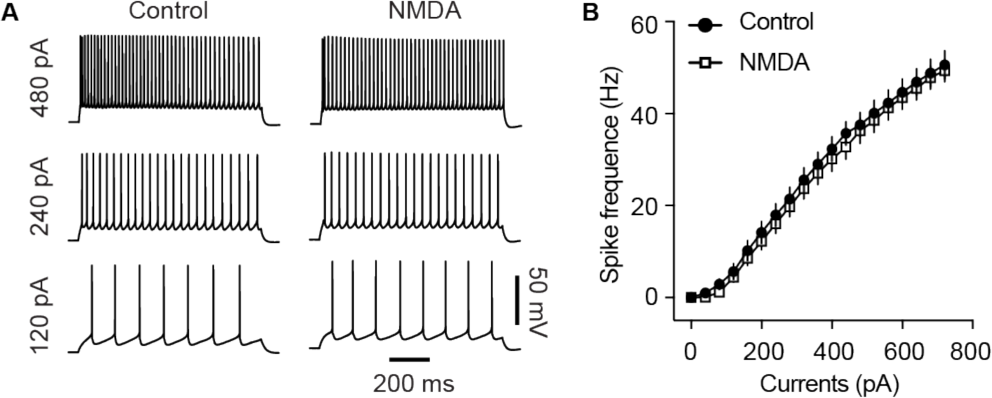
*Ex vivo* NMDA receptor activation does not affect cellular excitability of M1 PT neurons. **A**) Representative traces of action potentials of PT neurons from slices treated with normal ACSF (control) or ACSF containing NMDA (25 μM). **B**) Summarized data.

**Supplementary Figure 3.**
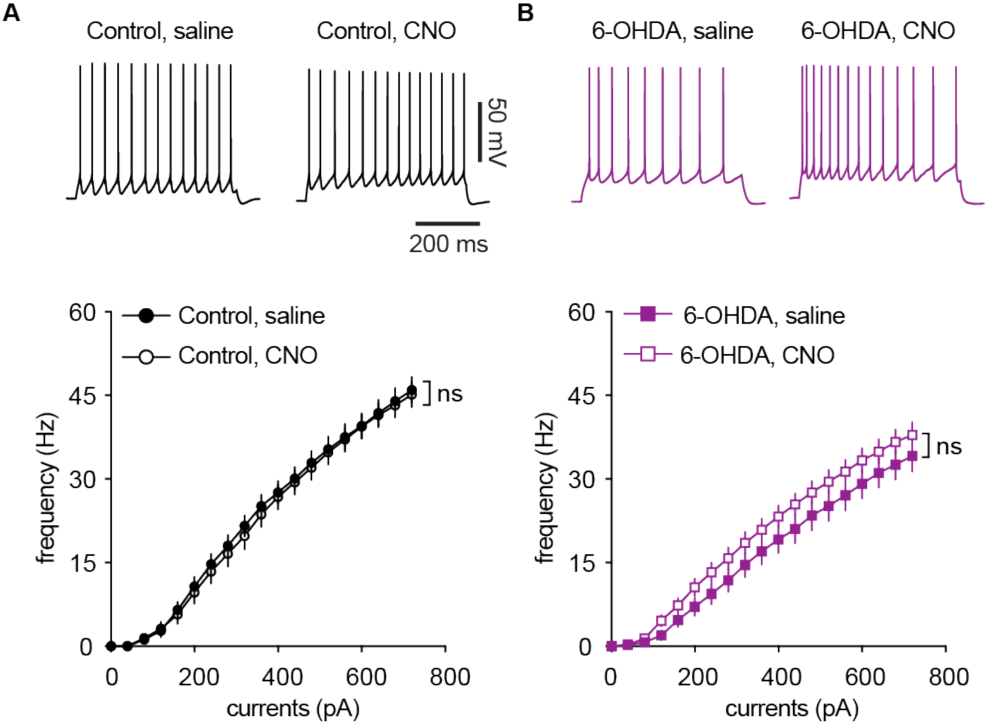
Chemogenetic suppression of basal ganglia output does not affect intrinsic excitability of M1 PT neurons. **A**) Representative traces of action potentials of PT neurons from saline- and CNO-injected control mice (top) and summarized results (bottom). **B**) Representative traces of action potentials of PT neurons from saline- and CNO-injected 6-OHDA mice (top) and summarized results (bottom).

## Notes

https://osf.io/m9dyw/?view_only=c96a43dbe04c4acf95c04c3cc7059ae5.

### Competing Interest Statement

The authors have declared no competing interest.

### Summary of Updates

per journal policy, the very original version o this work was re-posted.

## References cited

1. Aeed, F., Cermak, N., Schiller, J., and Schiller, Y. (2021). Intrinsic Disruption of the M1 Cortical Network in a Mouse Model of Parkinson’s Disease. Movement Disord 36, 1565–1577. https://doi.org/10.1002/mds.28538.

2. Albin, R.L., Young, A.B., and Penney, J.B. (1989). The functional anatomy of basal ganglia disorders. Trends Neurosci 12, 366–375. https://doi.org/10.1016/0166-2236(89)90074-x.

3. Biane, J.S., Takashima, Y., Scanziani, M., Conner, J.M., and Tuszynski, M.H. (2016). Thalamocortical Projections onto Behaviorally Relevant Neurons Exhibit Plasticity during Adult Motor Learning. Neuron 89, 1173–1179. https://doi.org/10.1016/j.neuron.2016.02.001.

4. Brazhnik, E., McCoy, A.J., Novikov, N., Hatch, C.E., and Walters, J.R. (2016). Ventral Medial Thalamic Nucleus Promotes Synchronization of Increased High Beta Oscillatory Activity in the Basal Ganglia–Thalamocortical Network of the Hemiparkinsonian Rat. J Neurosci 36, 4196–4208. https://doi.org/10.1523/jneurosci.3582-15.2016.

5. Brown, A.R., Hu, B., Antle, M.C., and Teskey, G.C. (2009). Neocortical movement representations are reduced and reorganized following bilateral intrastriatal 6-hydroxydopamine infusion and dopamine type-2 receptor antagonism. Exp Neurol 220, 162–170. https://doi.org/10.1016/j.expneurol.2009.08.015.

6. Bruno, R.M., and Sakmann, B. (2006). Cortex Is Driven by Weak but Synchronously Active Thalamocortical Synapses. Science 312, 1622–1627. https://doi.org/10.1126/science.1124593.

7. Chen, L., Daniels, S., Kim, Y., and Chu, H.-Y. (2021). Cell Type-Specific Decrease of the Intrinsic Excitability of Motor Cortical Pyramidal Neurons in Parkinsonism. J Neurosci 41, 5553–5565. https://doi.org/10.1523/jneurosci.2694-20.2021.

8. Chu, H.-Y., McIver, E.L., Kovaleski, R.F., Atherton, J.F., and Bevan, M.D. (2017). Loss of hyperdirect pathway cortico-subthalamic inputs following degeneration of midbrain dopamine neurons. Neuron 95, 1306–1318.e5. https://doi.org/10.1016/j.neuron.2017.08.038.

9. Cousineau, J., Lescouzères, L., Taupignon, A., Delgado-Zabalza, L., Valjent, E., Baufreton, J., and Bon-Jégo, M.L. (2020). Dopamine D2-like receptors modulate intrinsic properties and synaptic transmission of parvalbumin interneurons in the mouse primary motor cortex. Eneuro ENEURO.0081-20.2020. https://doi.org/10.1523/eneuro.0081-20.2020.

10. Galvan, A., and Wichmann, T. (2008). Pathophysiology of parkinsonism. Clin Neurophysiology Official J Int Fed Clin Neurophysiology 119, 1459–1474. https://doi.org/10.1016/j.clinph.2008.03.017.

11. Gaspar, P., Duyckaerts, C., Alvarez, C., Javoy-Agid, F., and Berger, B. (1991). Alterations of dopaminergic and noradrenergic innervations in motor cortex in Parkinson’s disease. Ann Neurol 30, 365–374. https://doi.org/10.1002/ana.410300308.

12. Georgopoulos, A.P., and Carpenter, A.F. (2015). Coding of movements in the motor cortex. Curr Opin Neurobiol 33, 34–39. https://doi.org/10.1016/j.conb.2015.01.012.

13. Goldberg, J.A., Boraud, T., Maraton, S., Haber, S.N., Vaadia, E., and Bergman, H. (2002). Enhanced synchrony among primary motor cortex neurons in the 1-Methyl-4-Phenyl-1,2,3,6-Tetrahydropyridine primate model of Parkinson’s disease. J Neurosci 22, 4639–4653. https://doi.org/10.1523/jneurosci.22-11-04639.2002.

14. Guo, J.-Z., Graves, A.R., Guo, W.W., Zheng, J., Lee, A., Rodríguez-González, J., Li, N., Macklin, J.J., Phillips, J.W., Mensh, B.D., et al. (2015a). Cortex commands the performance of skilled movement. Elife 4, e10774. https://doi.org/10.7554/elife.10774.

15. Guo, L., Xiong, H., Kim, J.-I., Wu, Y.-W., Lalchandani, R.R., Cui, Y., Shu, Y., Xu, T., and Ding, J.B. (2015b). Dynamic rewiring of neural circuits in the motor cortex in mouse models of Parkinson’s disease. Nat Neurosci 18, 1299–1309. https://doi.org/10.1038/nn.4082.

16. Hooks, B.M., Mao, T., Gutnisky, D.A., Yamawaki, N., Svoboda, K., and Shepherd, G.M.G. (2013). Organization of Cortical and Thalamic Input to Pyramidal Neurons in Mouse Motor Cortex. J Neurosci 33, 748–760. https://doi.org/10.1523/jneurosci.4338-12.2013.

17. Hyland, B.I., Seeger-Armbruster, S., Smither, R.A., and Parr-Brownlie, L.C. (2019). Altered recruitment of motor cortex neuronal activity during the grasping phase of skilled reaching in a chronic rat model of unilateral Parkinsonism. J Neurosci 39, 0720–19. https://doi.org/10.1523/jneurosci.0720-19.2019.

18. Kim, J., Kim, Y., Nakajima, R., Shin, A., Jeong, M., Park, A.H., Jeong, Y., Jo, S., Yang, S., Park, H., et al. (2017). Inhibitory Basal Ganglia Inputs Induce Excitatory Motor Signals in the Thalamus. Neuron 95, 1181–1196.e8. https://doi.org/10.1016/j.neuron.2017.08.028.

19. Kuramoto, E., Furuta, T., Nakamura, K.C., Unzai, T., Hioki, H., and Kaneko, T. (2009). Two Types of Thalamocortical Projections from the Motor Thalamic Nuclei of the Rat: A Single Neuron-Tracing Study Using Viral Vectors. Cereb Cortex 19, 2065–2077. https://doi.org/10.1093/cercor/bhn231.

20. Lafourcade, M., Goes, M.-S.H. van der, Vardalaki, D., Brown, N.J., Voigts, J., Yun, D.H., Kim, M.E., Ku, T., and Harnett, M.T. (2022). Differential dendritic integration of long-range inputs in association cortex via subcellular changes in synaptic AMPA-to-NMDA receptor ratio. Neuron https://doi.org/10.1016/j.neuron.2022.01.025.

21. McColgan, P., Joubert, J., Tabrizi, S.J., and Rees, G. (2020). The human motor cortex microcircuit: insights for neurodegenerative disease. Nat Rev Neurosci 21, 401–415. https://doi.org/10.1038/s41583-020-0315-1.

22. McGregor, M.M., and Nelson, A.B. (2019). Circuit mechanisms of Parkinson’s disease. Neuron 101, 1042–1056. https://doi.org/10.1016/j.neuron.2019.03.004.

23. McIver, E.L., Atherton, J.F., Chu, H.-Y., Cosgrove, K.E., Kondapalli, J., Wokosin, D., Surmeier, D.J., and Bevan, M.D. (2019). Maladaptive downregulation of autonomous subthalamic nucleus activity following the loss of midbrain dopamine neurons. Cell Reports 28, 992–1002.e4. https://doi.org/10.1016/j.celrep.2019.06.076.

24. Miller, O.H., Bruns, A., Ammar, I.B., Mueggler, T., and Hall, B.J. (2017). Synaptic Regulation of a Thalamocortical Circuit Controls Depression-Related Behavior. Cell Reports 20, 1867–1880. https://doi.org/10.1016/j.celrep.2017.08.002.

25. Oswal, A., Brown, P., and Litvak, V. (2013). Synchronized neural oscillations and the pathophysiology of Parkinson’s disease. Curr Opin Neurol 26, 662–670. https://doi.org/10.1097/wco.0000000000000034.

26. Pasquereau, B., and Turner, R.S. (2011). Primary motor cortex of the parkinsonian monkey: differential effects on the spontaneous activity of pyramidal tract-type neurons. Cereb Cortex 21, 1362–1378. https://doi.org/10.1093/cercor/bhq217.

27. Pasquereau, B., DeLong, M.R., and Turner, R.S. (2016). Primary motor cortex of the parkinsonian monkey: altered encoding of active movement. Brain 139, 127–143. https://doi.org/10.1093/brain/awv312.

28. Petreanu, L., Huber, D., Sobczyk, A., and Svoboda, K. (2007). Channelrhodopsin-2– assisted circuit mapping of long-range callosal projections. Nat Neurosci 10, 663–668. https://doi.org/10.1038/nn1891.

29. Plowman, E.K., Thomas, N.J., and Kleim, J.A. (2011). Striatal Dopamine Depletion Induces Forelimb Motor Impairments and Disrupts Forelimb Movement Representations within the Motor Cortex. J Park Dis 1, 93–100. https://doi.org/10.3233/jpd-2011-11017.

30. Sauerbrei, B.A., Guo, J.-Z., Cohen, J.D., Mischiati, M., Guo, W., Kabra, M., Verma, N., Mensh, B., Branson, K., and Hantman, A.W. (2020). Cortical pattern generation during dexterous movement is input-driven. Nature 577, 386–391. https://doi.org/10.1038/s41586-019-1869-9.

31. Shenoy, K.V., Sahani, M., and Churchland, M.M. (2012). Cortical Control of Arm Movements: A Dynamical Systems Perspective. Annu Rev Neurosci. 36, 337–359. https://doi.org/10.1146/annurev-neuro-062111-150509.

32. Turrigiano, G. (2011). Too Many Cooks? Intrinsic and Synaptic Homeostatic Mechanisms in Cortical Circuit Refinement. Annu Rev Neurosci 34, 89–103. https://doi.org/10.1146/annurev-neuro-060909-153238.

33. Viaro, R., Morari, M., and Franchi, G. (2011). Progressive motor cortex functional reorganization following 6-hydroxydopamine lesioning in rats. J Neurosci 31, 4544– 4554. https://doi.org/10.1523/jneurosci.5394-10.2011.

34. Villalba, R.M., Behnke, J.A., Pare, J.-F., and Smith, Y. (2021). Comparative Ultrastructural Analysis of Thalamocortical Innervation of the Primary Motor Cortex and Supplementary Motor Area in Control and MPTP-Treated Parkinsonian Monkeys. Cereb Cortex https://doi.org/10.1093/cercor/bhab020.

35. Villalobos, C.A., and Basso, M.A. (2022). Optogenetic activation of the inhibitory nigro-collicular circuit evokes contralateral orienting movements in mice. Cell Reports 39, 110699. https://doi.org/10.1016/j.celrep.2022.110699.

36. Vitrac, C., and Benoit-Marand, M. (2017). Monoaminergic Modulation of Motor Cortex Function. Front Neural Circuit 11, 72. https://doi.org/10.3389/fncir.2017.00072.

37. Wang, H., Stradtman, G.G., Wang, X.-J., and Gao, W.-J. (2008). A specialized NMDA receptor function in layer 5 recurrent microcircuitry of the adult rat prefrontal cortex. Proc National Acad Sci 105, 16791–16796. https://doi.org/10.1073/pnas.0804318105.

38. West, M.J. (1999). Stereological methods for estimating the total number of neurons and synapses: issues of precision and bias. Trends Neurosci 22, 51–61. https://doi.org/10.1016/s0166-2236(98)01362-9.

39. Wojcik, S.M., Rhee, J.S., Herzog, E., Sigler, A., Jahn, R., Takamori, S., Brose, N., and Rosenmund, C. (2004). An essential role for vesicular glutamate transporter 1 (VGLUT1) in postnatal development and control of quantal size. Proc National Acad Sci 101, 7158–7163. https://doi.org/10.1073/pnas.0401764101.

40. Xu, T., Yu, X., Perlik, A.J., Tobin, W.F., Zweig, J.A., Tennant, K., Jones, T., and Zuo, Y. (2009). Rapid formation and selective stabilization of synapses for enduring motor memories. Nature 462, 915–919. https://doi.org/10.1038/nature08389.

